# Pyramidal neuron morphogenesis requires a septin network that stabilizes filopodia and suppresses lamellipodia during neurite initiation

**DOI:** 10.1101/2022.06.19.496721

**Authors:** Megan R. Radler, Xiaonan Liu, Megan Peng, Brenna Doyle, Kazuhito Toyo-Oka, Elias T. Spiliotis

## Abstract

Pyramidal neurons are the major cell type of the forebrain, consisting of a pyramidally shaped soma with axonal and apicobasal dendritic processes. It is poorly understood how the neuronal soma morphs from a sphere to pyramid, while generating neurites of the proper shape and orientation. Here, we discovered that the spherical somata of immature neurite-less neurons possess a circumferential wreath-like network of septin filaments, which promotes myosin II localization and suppresses Arp2/3 activity at the base of filopodial actin bundles. The septin network facilitates neurite formation by stabilizing nascent filopodia, which mature to neurites, and concomitantly maintains a consolidated soma by suppressing the extension of lamellipodia. We show that this septin function is critical for the morphogenesis and spatial orientation of pyramidal somata and their neurites in vitro and in vivo. Therefore, the somatic septin cytoskeleton provides a key morphogenetic mechanism for neuritogenesis and the development of pyramidal neurons.

**Highlights:**

- A septin wreath-like network controls the shape of neuronal somata and nascent neurites
- Septins promote and suppress filopodial and lamellipodial protrusions, respectively
- Septins scaffold myosin II and exclude Arp2/3 at the base of filopodial actin
- Development of pyramidally shaped neurons requires septins in vitro and in vivo

**eTOC Summary:** Radler et al report a new morphogenetic mechanism in the development of pyramidal neurons, which is mediated by a septin wreath-like cytoskeleton in the soma of immature spherical neurons. The septin network stabilizes somatic filopodia and suppresses lamellipodia by differentially controlling the localization of myosin II and Arp2/3.

## Introduction

In the forebrain, excitatory neurons have pyramidally shaped cell bodies (somata) with axonal and apicobasal dendritic processes, which originate from the apex and base of their pyramidal shapes (Mohan, et al., 2015; Spruston, 2008). Understanding how pyramidal neurons generate and maintain their specialized morphology is imperative for tackling neurodevelopmental and neurodegenerative disorders, and devising neuroregenerative strategies (Liu and Jan, 2020; Prem, et al., 2020; Copf, 2016; Kulkarni and Firestein, 2012).

Neurites, the precursors of axons and dendrites, originate from the membrane protrusions of the neuronal soma (Miller and Suter, 2018; Sainath and Gallo, 2015; da Silva and Dotti, 2002). In the developing brain, neurites develop from the leading and trailing processes of migrating neurons (Takano, et al., 2019; Barnes and Polleux, 2009). However, neurite outgrowth also occurs prior to migration, resembling the stochastic formation and differentiation of multiple neurites in cultured neurons (Namba, et al., 2014; Hatanaka and Yamauchi, 2013; Dotti, et al., 1988). In vitro, morphogenesis of pyramidal neurons initiates from spherically shaped cells, which intrinsically break their radial symmetry and extend neurites that differentiate into axonal and apicobasal-like dendritic processes (Banker, 2018; da Silva and Dotti, 2002).

Neurite formation is mechanistically executed by the actin and microtubule cytoskeleton (Miller and Suter, 2018; Sainath and Gallo, 2015; Flynn, 2013). Polymerization of branched actin, which is nucleated by the Arp2/3 machinery of actin assembly, extends lamellipodia protrusions from the peripheral lamellae of spherical neurons (Gautreau, et al., 2022; Sainath and Gallo, 2015; Svitkina, 2013). Neurites develop from filopodia which form by reorganization of the lamellipodial actin network and/or actin polymerization at microdomains of the cortical membrane (Wit and Hiesinger, 2022; Svitkina, 2013; Saengsawang, et al., 2012; Guerrier, et al., 2009; Dent, et al., 2007; Kwiatkowski, et al., 2007). Neurites can also emerge from microtubule-driven protrusions, or the collapse and consolidation of lamellipodia into cylindrical structures, which is facilitated by microtubule entry and concomitant suppression of actin-based protrusion along the neurite shaft (Rao and Baas, 2018; Lu, et al., 2013; Flynn, et al., 2012; Mingorance-Le Meur and O’Connor, 2009; Dehmelt, et al., 2003). Despite advances in our mechanistic understanding of neurite formation, it is unknown how the neuronal soma develops from a sphere into a pyramid, while generating neurites of the right shape and orientation. This morphogenetic development requires spatial regulation and coordination of the lamellipodial and filopodial protrusion of the soma, but the underlying mechanisms are little understood.

Septins are GTP-binding proteins, which assemble into non-polar filamentous oligomers and polymers that associate with membranes and the cytoskeleton (Spiliotis and Nakos, 2021; Woods and Gladfelter, 2021). Septins have evolutionarily conserved roles in cellular morphogenesis and the spatial organization of membrane and cytoskeletal proteins (Spiliotis and Nakos, 2021; Caudron and Barral, 2009). In neurons, septins have been functionally linked to axodendritic growth and branching, dendritic spine maturation and polarized membrane traffic (Radler and Spiliotis, 2022; Ageta-Ishihara and Kinoshita, 2021). Septin 7 (Sept7), a ubiquitous and obligate subunit of heteromeric septin complexes, was notably found to provide a spatial memory for the post-mitotic reemergence of neurites in neural crest cells (Boubakar, et al., 2017). How Sept7 controls neuritogenesis, however, is unknown, and septin functions in the early stages of neuronal morphogenesis have not been explored. Here, we have discovered that the soma of neurite-less spherical neurons contains a novel network of Sept7-containing filaments, which biases membrane protrusive activity toward filopodia by promoting and suppressing myosin II and Arp2/3, respectively. Through this mechanism, Sept7 promotes the formation of properly shaped neurites while maintaining a compact soma, which is critical for the formation of a pyramidally shaped soma with axodendritic processes of proper morphology and orientation.

## Results

### Spherical stage 1 neurons possess a circumferential septin wreath-like network that demarcates a zone of myosin II enrichment and Arp2/3 diminution

Septins have been implicated in neuritogenesis, but it is unknown how they function in neurite formation (Falk, et al., 2019). We therefore set out to examine septin localization and function in the early stages of neurite formation in pyramidal neurons. We focused on septin 7 (Sept7), as a ubiquitous subunit of mammalian complexes, in embryonic stage 1 (DIV0) rat hippocampal (E18) and mouse cortical (E15) neurons. Using super-resolution confocal and structured illumination microscopy (SIM), we observed a circumferential wreath-like network of Sept7 filaments which was positioned at the interface of the central microtubule meshwork and the actin network of the peripheral lamellae (Figure 1A and S1). Consistent with previous reports of Sept5/11/7 complexes in the soma of hippocampal neurons (Xie, et al., 2007), we found that Sept5 and Sept11 had a similar localization and appearance (Figure S1A-B). In mouse cortical neurons, Sept5 and Sept11 also colocalized with the Sept7 network (Figure S1C-E). The circumferential septin network consisted of linear and curved filaments of variable lengths and thicknesses (Figure 1A). While devoid of a regular pattern, the septin network contained an array of spoke-like curvilinear fibers, which are positioned orthogonally to the inner perimeter of the cell body (Figure 1A, arrowheads). In contrast to hitherto known mammalian septin networks, which are largely integrated with the microtubule and actin cytoskeleton, the circumferential septin was positionally and structurally distinct (Figure 1A). We observed, however, regions of Sept7 colocalization with the proximal ends of filipodial actin bundles (Figure 1A, arrows). Sept7 accumulation at the base of filopodial bundles was also observed in neurons, which had broken spherical symmetry (Figure 1E). Sept7 was concentrated along proximal segments of filopodial actin bundles, which were either fully embedded in the veils of peripheral lamellae of spherical neurons (Figure 1A-B) or protruded in a neurite-like manner from the soma of asymmetrically shaped neurons (Figure 1E).

**Figure 1.**
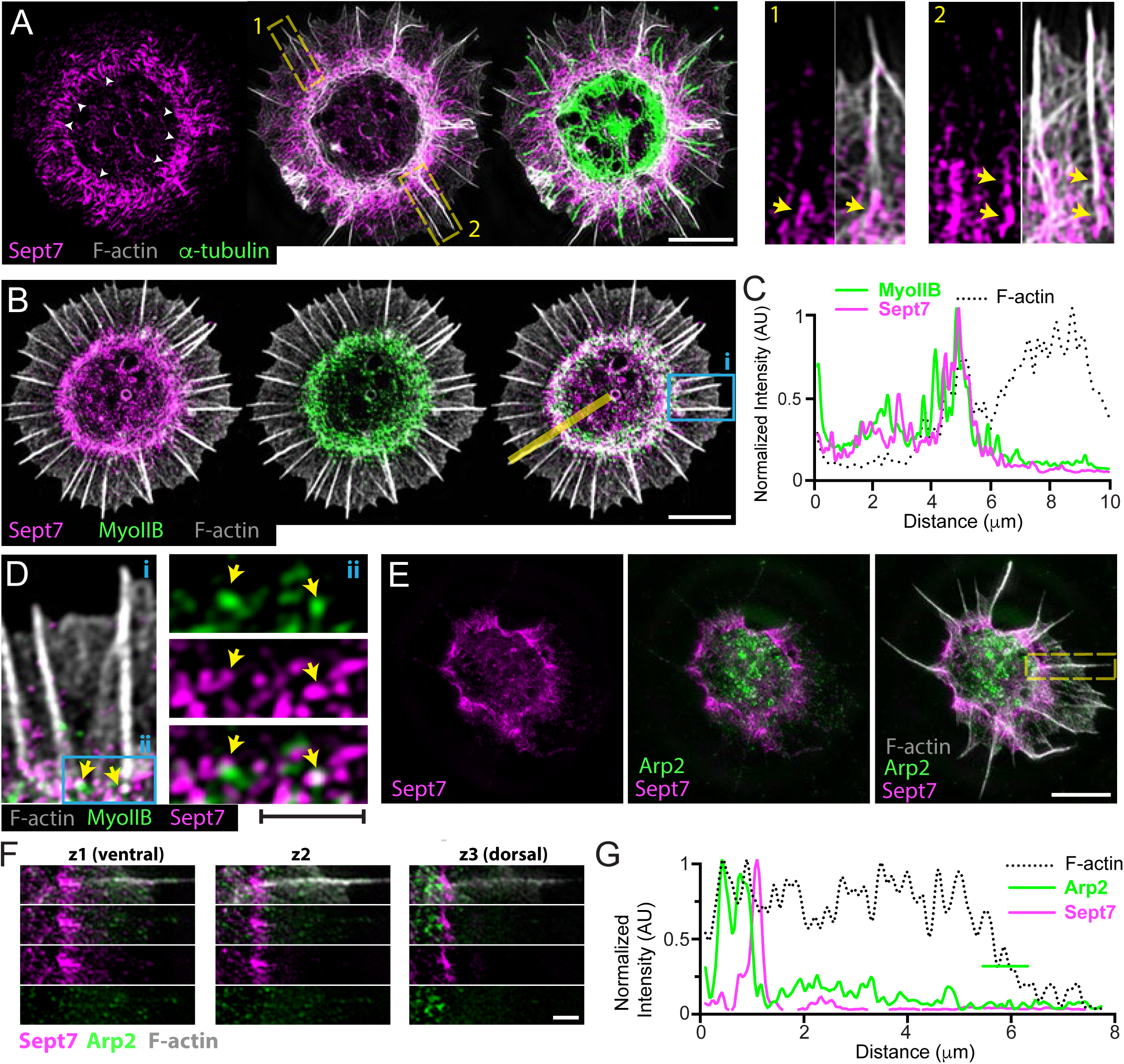
A circumferential septin wreath-like network demarcates a zone of myosin II enrichment and Arp2 diminution at the base of filopodial actin bundles. (A) Structured illumination microscopy (SIM) images of rat hippocampal neuron (DIV0) stained for Sept7, F-actin (phalloidin) and α-tubulin. White arrowheads point to spoke-like curve linear fibers with an orthogonal orientation to the inner perimeter of the soma. Insets show in higher magnification the localization of Sept7 (arrows) at the base of filopodial actin bundles. Scale bar, 5 μm. (B-C) Super-resolution SIM images (B) of rat hippocampal neuron (DIV0) stained for Sept7, myosin IIB, and F-actin (phalloidin). Intensity profile plot (C) shows the fluorescence intensity levels of Sept7, myosin IIB and F-actin across the yellow line from the center to the periphery of the neuron. Scale bar, 5 μm. (D) A selected region (i) from panel B is shown in higher magnification, and outlined area (ii) shows myosin IIB and Sept7 overlap at the base of filopodial actin bundles. Scale bar, 1 μm. (E-G) SIM image of rat hippocampal neuron (DIV0) stained for Arp2, Sept7 and F-actin (phalloidin; E). Optical sections from the ventral, medial and dorsal sides of an outlined filopodial actin bundle are shown in higher magnification (F). Fluorescence intensity plot (G) shows diminution of Arp2 fluorescence at the region of Sept7 localization at the base of filopodial actin. Line-scan quantification was performed in the optical section (z3) with the highest enrichment of Arp2 at the base of the outlined filopodial actin bundle. Scale bars, 5 μm (E) and 1 μm (F).

We next sought to determine whether the circumferential septin network demarcated or colocalized with any actin-binding proteins. We stained for the non-muscle myosin II, which is a known septin-interacting protein, and Arp2/3 which nucleates the actin filaments of the peripheral lamellae. Strikingly, myosin IIB localized as a circumferential band of short fibers and puncta, which overlapped with the septin network (Figure 1B). The myosin IIB and septin networks were interwoven, and myosin IIB overlapped and colocalized with Sept7 at the base of filopodial actin bundles (Figure 1C-D). In contrast to myosin II, Arp2 did not have a circumferential pattern. Patches of Arp2 puncta localized to the posterior of the septin network and along the central area of the soma (Figure 1E). In the circumferential septin network, Arp2 puncta were sparse and largely absent from or to the posterior of Sept7 accumulations at the base of filopodial actin bundles (Figure 1F-G). Thus, prior to neurite formation, immature neurons contain a circumferential septin network, which demarcates a zone of myosin II enrichment and Arp2 diminution at the base of the filopodial actin bundles of peripheral lamellae.

### Sept7-depleted neurons have an enlarged soma with hyperextended lamellae and defective neurite outgrowth

To test whether the circumferential septin network has a role in neurite initiation and the morphogenesis of pyramidal neurons, we knocked down Sept7 expression by transfecting rat hippocampal neurons (DIV1) with Sept7-specific shRNAs for 48 h (Figure S2A-B). Sept7 depletion resulted in a dramatic enlargement of the soma without altering the size of the nucleus (Figure 2A-C, and S2C-D). The soma of Sept7-depleted neurons was characterized by hyper-extended spread-out lamellae, which contained multiple lamellipodia, indicative of unrestrained lamellipodial activity (Figure 2A). This phenotype was accompanied by aberrant neurite outgrowth. Neurites emerged mainly from the edges of lamellipodia (Figure 2A; red colored neurites), which stretched from the soma, instead of originating directly from the narrowing edges of a compact soma, which is the main mode of neurite outgrowth in control cells (Figure 2A, D). The lamellipodia-derived neurites of Sept7-depleted neurons were thinner than the somatic neurites of control cells, resulting in an overall decrease in the mean neurite width, which was measured at 5 μm from from the point of origin (Figure 2E). Although neurite length was not affected (Figure S2E), the total number of neurites per cell and the number of neurite branches increased in Sept7-depleted cells, which was consistent with an enhanced protrusive activity (Figure 2F-G). These phenotypes were not due to non-specific off-target effects as they were rescued upon expression of GFP-Sept7, which was not targeted by the shRNA against the 3’ untranslated region of Sept7 (Figure S2C-D, and S2F-I).

**Figure 2.**
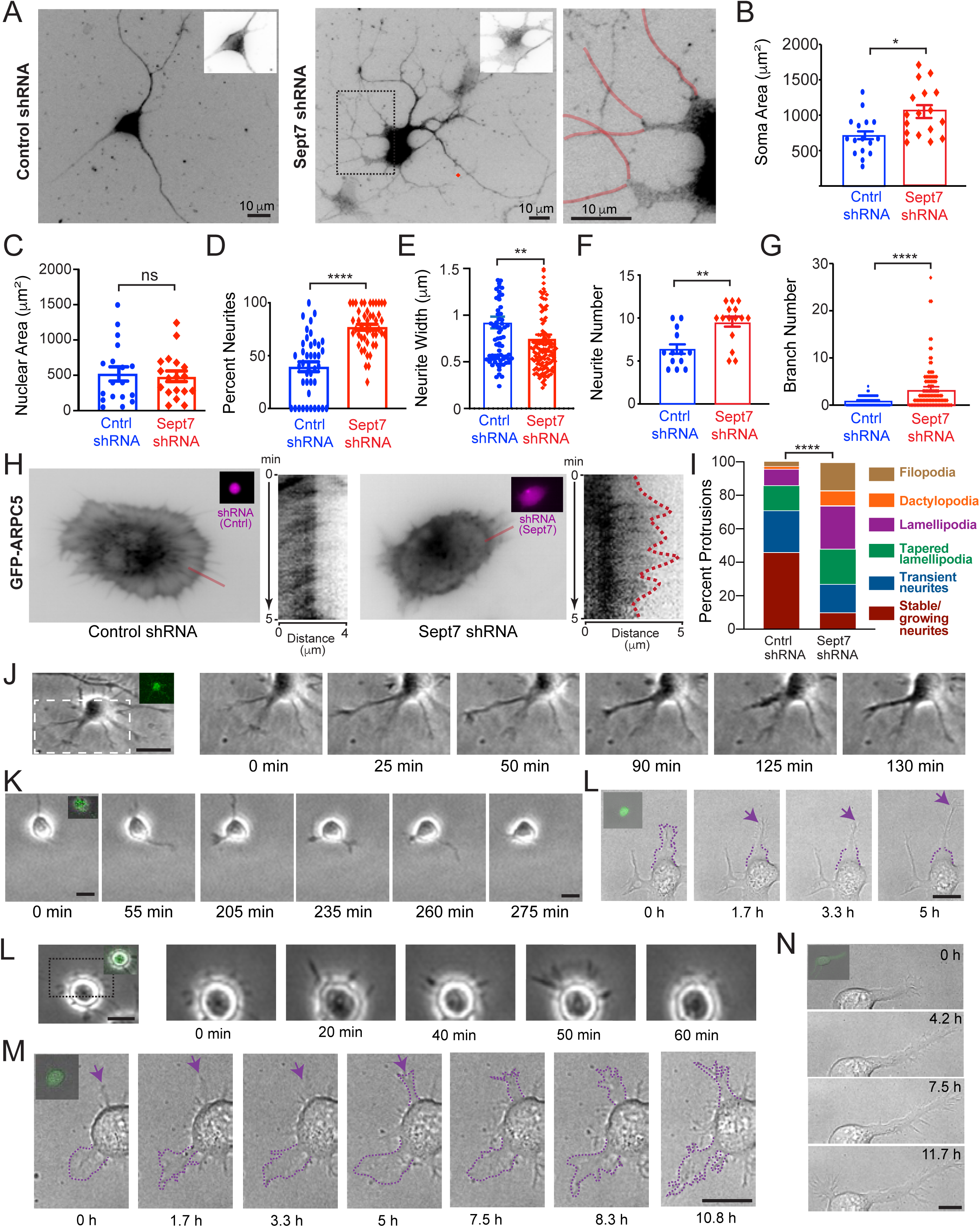
Neurite morphology requires septin-dependent suppression of the lamellipodial protrusive activity of the soma. (A) Rat hippocampal neurons (DIV1) were transfected with plasmids that co-express GFP and shRNAs (Sept7 3’-UTR or scrambled control) for 48 h, and stained for endogenous Sept7 (inset). An outlined region with neurites originating from hyperextended lamellipodia (pseudo-colored in red) is shown in higher magnification. Scale bars, 10 μm. (B-C) Bar graphs show mean (± SEM) surface area of the soma (B) and the nucleus (C) of rat hippocampal neurons (n = 18) transfected with control and Sept7 shRNAs for 48 h. Data were analyzed with the Mann-Whitney U test. (D) Quantification of percent of neurites per neuron (n = 40-46 neurons) that emerge from extended lamellipodial protrusions of the soma. Data were analyzed with a student’s test. (E-G) Mean (± SEM) neurite width (E; n = 80-137), neurite number per neuron (D; n = 13-15), and neurite branches per neuron (F; n = 82-115). Data were analyzed from 13-16 neurons that were transfected with scrambled control or Sept7 shRNAs for 48 h, and were analyzed with the Mann-Whitney U test. (H) Rat hippocampal neurons (DIV1) were transfected with shRNAs for 48 h, trypsinized and re-plated for 1 h before imaging. Images show the sum projection of the frames of a 5 minute time-lapse movie of GFP-ARPC5, which was acquired with total internal reflection fluorescence (TIRF) microscopy. Insets show the mCherry fluorescence that reports on expression of control or Sept7 shRNAs. Red lines outline the linear region of the kymographs that demonstrate the dynamics of GFP-ARPC5 in control and Sept7 knock-down neurons. (I) Rat hippocampal neurons (DIV1) were transfected with shRNAs for 48 h, trypsinized, re-plated and imaged live in 5 minute intervals with phase contrast or differential interference contrast (DIC) microscopy. Stacked bar graph shows the percentage of the protrusions of the soma, which were categorized into: i) stably growing or transient (growing/retracting) neurites, which originate from the soma, iii) neurites that emerge from the tapering ends of lamellipodia or are connected to lamellipodia (tapered lamellipodia), iv) dactylopodia-like neurites, and v) filopodia. The morphology of 93-125 neurites from 16-18 neurons was analyzed, and statistical significance was calculated with a chi squared test. ****, p < 0.0001 (J-L) Phase-contrast (J, K) and DIC (L) still-frame images from overnight movies of living hippocampal neurons that were re-plated after transfection with scrambled control shRNAs for 48 h. Insets show GFP as a reporter of shRNA expression. Lamellipodial protrusions of the soma are outlined with a dotted line and arrows point to the formation of a neurite from the tapering end of a lamellipodium. Scale bars, 10 μm. (L-N) Phase-contrast (L) and DIC (M, N) images from overnight time-lapse imaging of living hippocampal neurons that were re-plated after transfection with Sept7 shRNAs for 48 h. Insets show GFP as a reporter of shRNA expression. Scale bars, 10 μm. Statistics. *p<0.05, **p<0.01, ***p<0.001, ****p<0.0001, ns: non-significant

To better resolve how neurite formation is impacted by Sept7 depletion, we imaged neurons that were transfected with shRNAs for 48 h, and then were detached and replated to reinitiate neuritogenesis. After one hour of replating, we first imaged membrane dynamics in living neurons expressing the Arp2/3 complex marker GFP-ARPC5. A cumulative projection of time-lapse frames and kymographs of cell edge dynamics showed that Sept7-depleted neurons lack well-defined lamellipodia and filopodia with retrograde actin flow (Figure 2H, Video S1). In contrast, cell edges consisted of undulated and ruffling lamellipodia, undergoing repeated bouts of extension (Figure 2H, Video S2)

Using phase contrast or differential interference contrast (DIC) time-lapse microscopy, we then imaged living neurons overnight to capture the generation and dynamics of nascent neurites. Consistent with previous studies, neurites originated from filopodia shaped protrusions, which emerge directly from the soma, and either remained stable with an overall growth (Figure 2J, Video S3) or underwent bouts of outgrowth and complete retraction (transient neurites; Figure 2K, Video S4). In addition, a number of neurites formed from the tapering ends of lamellipodial protrusions (Figure 2L). Quantifications showed that in control neurons, 71% of nascent neurites underwent stable growth or grew and shrank (Figure 2I). Strikingly, this percentage decreased dramatically to 27% of total protrusions in Sept7 knock-down (Figure 2I). The majority of Sept7-depleted neurons developed lamellipodia protrusions, which did not resolve to neurites (26%) or formed neurites form tapering edges (21%; Figure 2I). Sept7-depleted neurons developed short (< 1 μm) and transient filopodia, which failed to elongate into neurites (Figure 2L; Video S5) and/or converted to wider lamellipodia-like protrusions (Figure 2M, arrow; Video S6). Taken together with the presence of elongated protrusions with a lamellipodia-like width that resembled dactylopodia (Figure 2M-N; Video S7), the invasive lamellipodia-related protrusions of endothelial cells (Figueiredo, et al., 2021), these phenotypes indicated an amplification of lamellipodial activity in the somata of Sept7-depleted neurons. Therefore, the circumferential Sep7-containing network of septins provides a key function in balancing the activity of filopodial and lamellipodial protrusions for the formation of neurites of the right shape and origin.

### The circumferential septin cytoskeleton promotes neurite formation by promoting myosin IIB localization and Arp2/3 exclusion at the base of filopodial actin bundles

Given that the circumferential septin network demarcates a zone of myosin II enrichment and Arp2/3 diminution at the base of the filopodial actin bundles of the soma (Figure 1), we hypothesized that it promotes neurite formation by spatially controlling myosin II and Arp2/3. We predicted that Sept7 knock-down may decrease myosin II localization, which would reduce retrograde actin flow and filopodia stability, and concomitantly may increase Arp2/3 levels, which would result in engorgement of filopodia with branched actin and their widening into lamellipodia- or dactylopodia-like protrusions (Alieva, et al., 2019; Yang, et al., 2012; Medeiros, et al., 2006; Lin, et al., 1996).

Using super-resolution SIM, we examined myosin IIB and Arp2 localization in the lamellae of neurons, which were re-plated after a 48 h treatment with shRNAs (DIV3) to reinitiate neurite formation. Quantification of myosin IIB at the proximal end of filopodial actin bundles showed a reduction in the percentage of myosin IIB-positive filopodia in Sept7-depleted neurons (Figure 3A-B). In contrast to this reduction of myosin IIB, Arp2 levels were visibly elevated at the base of filopodial actin bundles (Figure 3C). In control neurons, Arp2 was predominately concentrated at lamellipodial protrusions (Figure 3C, arrowhead), and exhibited a sparse punctate localization at the basal regions and along the length of filopodia (Figure 3C, arrows). In Sept7 knock-down neurons, however, Arp2 accumulated at the proximal segments of filopodia, and was more abundantly present in filopodia (Figure 3C-D). This Arp2 enrichment was evident not only at filopodia that originate directly from the soma (Figure 2C) but also in dactylopodia-like protrusions (Figure 2D, region 2), and filopodia that extend from the edges of lamellipodial protrusions (Figure 2D, region 1) resembling the aberrant neurites of the enlarged somata (Figure 2A).

**Figure 3.**
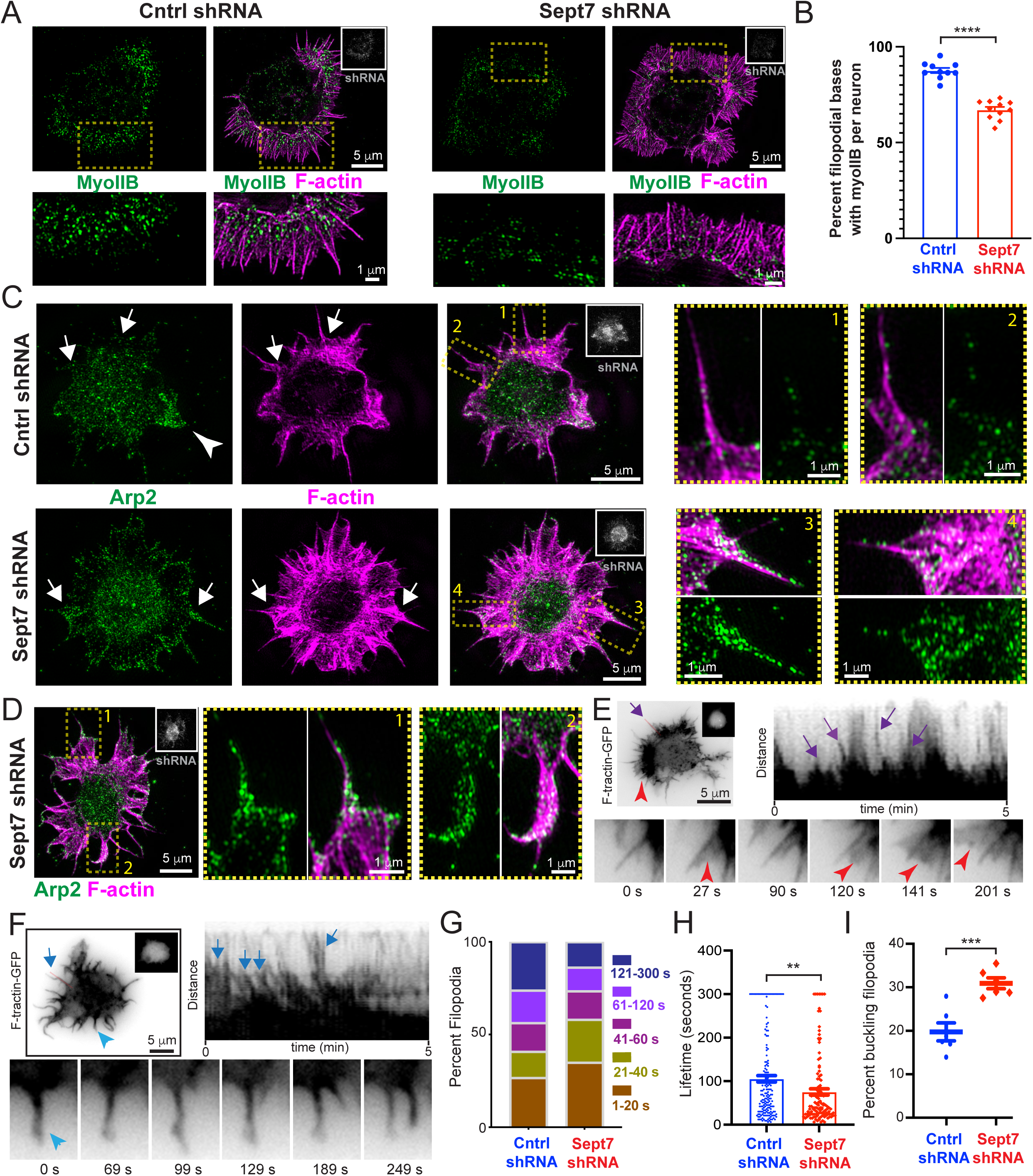
Sept7 biases somatic protrusions toward filopodia by controlling myosin II and Arp2 levels at the base of filopodial actin bundles. (A) SIM images of rat hippocampal neurons, which were transfected at DIV0 with plasmids that co-express GFP (inset) and scrambled control or Sept7 shRNAs for 48 h, trypsinized and re-plated for 2 h (scale bar, 5 μm). Outlined areas are shown in higher magnification (scale bar, 1μm) (B) Bar graph shows mean (± SEM) percentage of filopodial actin bundles (n > 40) per neuron with myosin IIB at their proximal basal ends. Filopodial actin bundles were analyzed from 10-11 neurons and statistical significance was derived using the unpaired t-test. (C-D) SIM images of rat hippocampal neurons (DIV0), which were transfected with plasmids that co-express GFP (inset) and scrambled control (C) or Sept7 shRNAs (C, D) for 48 h, trypsinized and re-plated for 2 h. Outlined areas are shown in higher magnification. Arrows point to the base of filopodial protrusions with paucity and enrichment of Arp2 in control and Sept7-depletd neurons, respectively. Arrowhead points to a lamellipodial protrusion, which is enriched with Arp2. Examples of Arp2 accumulation in filopodia (D), which extend from hyper-extended lamella, and dactylopodia-like protrusions are shown in higher magnification. (E-F) Still frames and kymographs from time-lapse imaging of F-tractin-GFP with TIRF microscopy in hippocampal neurons transfected with Sept7 shRNAs (E) and control scrambled shRNAs (F). Inset images (grayscale) show the mCherry that co-expressed with shRNAs. Magenta arrows point to transient filopodia and red arrowheads point to lamellipodial protrusions that extend over filopodia (E). Blue arrows point to filopodia that persist over time. Red lines show the regions from which kymographs were generated. (G) Stacked bar graph shows the percentage of filopodia (n = 106-172) with lifetimes between 1-20, 21-40, 41-60, 61-120 and 121-300 seconds, which were quantified from 5 minute-long time-lapse movies of F-tractin-GFP in rat hippocampal neurons transfected with scrambled control and Sept7 shRNAs. Quantifications were performed in five neurons. (H-I) Bar graphs show the mean (± SEM) lifetime of somatic filopodia (H; n = 106-172) and mean (± SEM) percentage of buckling filopodia per neuronal soma (I; n = 5). Quantifications were performed in fiver neurons from each condition, which co-expressed F-tractin-GFP and shRNAs, and imaged for 5 minutes by time-lapse TIRF microscopy. Data were analyzed with the Mann-Whitney U test (H) and unpaired t-test (I). Statistics. **p<0.01, ***p<0.001, ****p<0.0001

We next used time-lapse TIRF imaging of living neurons, which expressed shRNAs and F-tractin-EGFP, to analyze the dynamics of actin-based protrusions and determine how they are impacted by the effects of Sept7 depletion on myosin IIB and Arp2. Sept7-depleted neurons were characterized by filopodia, which overlapped with and/or were engulfed by veils of dynamic lamellipodia (Figure 3E, Video S8), while control neurons had well-defined and persistent filopodia (Figure 3F; Video S9). Kymograph analysis showed that the filopodia of Sept7-depleted neurons were more transient (Figure 2E-F, magenta vs blue arrows) and often dissipated in the wake of extending lamellipodia (Figure 2E, red arrowhead). Quantification of filopodia life-times within a 5-minute period of imaging revealed that Sept7 knock-down reduced the percentage of long-lived filopodia (61-120 s, 121-300 s) and increased filopodia with shorter lifetimes (1-20 s, 21-40 s; Figure 3G). Consistent with an overall reduction in the stability of filopodia, the mean life-time of filopodia decreased and a higher percentage of filopodia bent or buckled (Figure 3H-I). Because myosin II association with the base of filopodia augments their stability (Alieva, et al., 2019), these phenotypes were in agreement with myosin IIB reduction and Arp2/3 increase, which enhances lamellipodia activity at the edge of the neuronal soma (Gautreau, et al., 2022; Yang, et al., 2012; Lin and Forscher, 1995).

We reasoned that if the soma and neurite phenotypes (Figure 2) were indeed due to elevated Arp2/3 and suppressed myosin II activities, they might be reversible by inhibiting Arp2/3-mediated actin polymerization or boosting myosin II contractility. We first targeted Arp2/3 activity by treating Sept7-depleted neurons for 24 h with low concentrations (1 μm) of the Arp2/3 inhibitor CK666 (Hetrick, et al., 2013). CK666 reversed the enlargement of the soma, the surface area of which was reduced to the levels of control neurons (Figure 4A-B). In addition, CK666 rescued neurite morphology as well as neurite number and branching (Figure 4C-E). Neurites emerged directly from a compact soma rather than the edges of hyperextend lamellipodia, and had similar levels of arborization with control neurons. We also targeted Arp2/3 activity with a non-pharmacological approach using a dominant-negative cortactin construct (DN-cortactin-GFP), which consists of the N-terminal Arp3-binding domain of cortactin (Weed, et al., 2000). This construct has been previously shown to promote neurite consolidation by suppressing protrusive activity along the shaft of nascent neurites (Mingorance-Le Meur and O’Connor, 2009). Similar to CK666, DN-cortactin rescued the soma and neurite phenotypes of Sept7-depleted neurons (Figure 4F-J).

**Figure 4.**
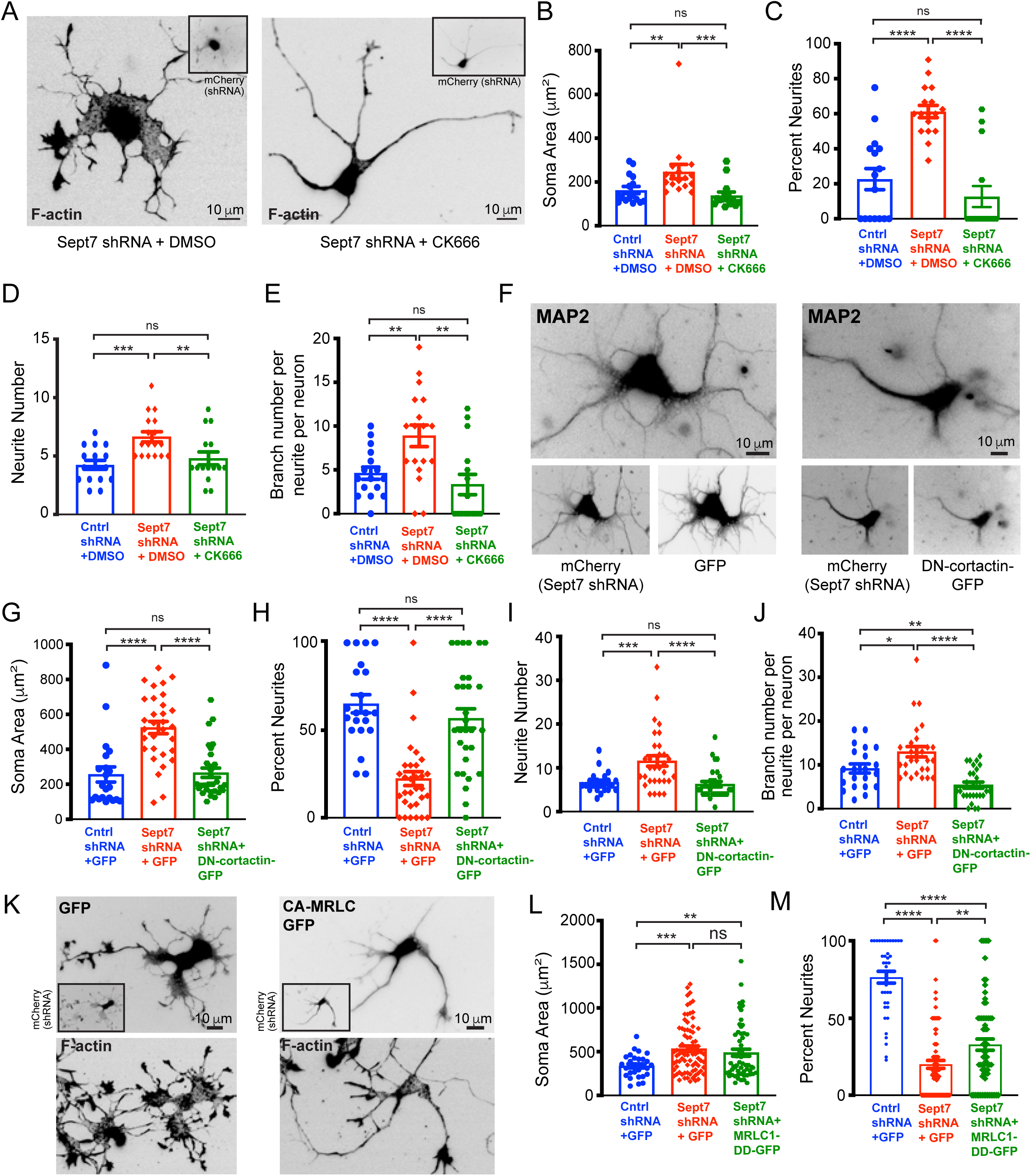
Inhibition of Arp2/3 activity rescues the morphology of the soma and neurites of Sept7-depleted hippocampal neurons. (A) Images show filamentous actin (phalloidin) in rat hippocampal neurons, which were transfected with plasmids that co-express mCherry (inset) and shRNAs for 48 h, and treated with CK666 (1 μM) for 24 h. (B-E) Bar graphs show the mean (± SEM) surface area of the soma (B), mean (± SEM) percentage of neurites per neuron which originate from hyperextended lamellipodia (C), mean (± SEM) number of neurites per neuron (D), and mean (± SEM) number of total neurite branches per neuron (E). All quantifications were performed in 15-17 rat hippocampal neurons, which were transfected with shRNAs on DIV1 for 48 h, and subsequently treated with CK666 (1 μM) for 24 h. Data were analyzed with the Mann-Whitney U test. (F) Images show MAP2 in rat hippocampal neurons, which were co-transfected with a plasmid that co-expresses mCherry and Sept7 shRNA, and GFP or GFP-tagged dominant negative cortactin (DN-cortactin-GFP). (G-J) Bar graphs show the mean (± SEM) surface area of the soma (G), mean (± SEM) percentage of neurites per neuron which originate directly from the soma (H), mean (± SEM) number of neurites per neuron (I), and mean (± SEM) number of neurite branches per neuron (J). Quantifications were performed in neurons that expressed control scrambled shRNAs and GFP (n = 21), Sept7 shRNAs and GFP (n = 31), Sept7 shRNAs and DN-cortactin-GFP (n = 30). Data were analyzed with Mann-Whitney U test. (K) Images show rat hippocampal neurons that express GFP or GFP-tagged constitutively active myosin II regulatory light chain (MRLC1-DD-GFP), and stained with phalloidin. Neurons were co-transfected with a plasmid that co-expresses mCherry and Sept7 shRNA, and GFP or GFP-tagged constitutively active myosin RLC (MRLC1-DD-GFP). (L-M) Bar graphs show the mean (± SEM) surface area of the soma (L), and mean (± SEM) percentage of neurites per neuron (M) that originated directly from the soma. Quantifications were performed in rat hippocampal neurons that were transfected with control scrambled shRNA and GFP (n = 30-39), Sept7 shRNA and GFP (n = 81-83) and Sept7 shRNA and MRLC1-DD-GFP (n = 59-67). Data were analyzed with the Mann-Whitney U test. Statistics. *p<0.05, **p<0.01, ***p<0.001, ****p<0.0001, ns: non-significant

We then tested if expression of the constitutively active di-phosphomimetic myosin regulatory light chain (MRLC1-DD-GFP) could rescue the phenotypes of Sept7 knock-down by enhancing myosin II contractility. MRLC1-DD did not reverse the hyperextended lamellae of the soma (Figure 4K-L), and the number of aberrant neurites that originate from the edges of spread out lamellipodia was incrementally reduced. The percentage of neurites that emerge from a compact soma increased from 20% to 38%, but was not rescued to the percentage (76%) of control neurons (Figure 4M). Neurite numbers and branching, however, were reduced to control levels (Figure S2J-K). These data indicated that MRLC1-DD expression was not sufficient to reverse the growing lamellipodia of the soma, which is likely due to the reduction of myosin II heavy chains from the circumferential lamellae of the soma, but it was more effective in other regions of neurite growth where myosin II heavy chains were more abundant.

Collectively, these data indicate that during neuritogenesis, the circumferential septin cytoskeleton minimizes lamellipodial formation by Arp2/3-driven polymerization, and promotes the stability of filopodia by scaffolding myosin II at the base of their actin bundles. Thus, septins promote the generation and maturation of filopodia into neurites from a consolidated compact soma.

### Septin function in early neuritogenesis is required for the development of a pyramidal soma with proper axodendritic orientation in vitro and in vivo

Our results reveal that early neurite formation requires regulation of the balance of filopodial and lamellipodial protrusions by a somatic network of septins. However, is this regulation critical for the morphogenesis of a pyramidally shaped neuron with an apicobasal dendritic tree? To answer this question, we examined the morphology of neurons that were allowed to develop for nine days after transfection with shRNAs (Figure 5A). We first analyzed the shape of the soma. In control neurons, ∼90% of somata were either triangular or quadrilateral, which are the predominate two-dimensional shapes of a pyramidal soma (Figure 5B). By contrast, only ∼50% of somata had a triangular or quadrilateral in Sept7-depleted neurons (Figure 5B). The rest of the neurons had pentagonal and other irregular multi-angular shapes (Figure 5B). Consistent with this phenotype, the surface area and complexity factor of the soma (ratio of perimeter length to surface area) were higher in Sept7-depleted neurons (Figure 5C-D). Hence, the enlargement and spreading of Sept7-depleted neuronal somata, which takes place in the early stage of neurite formation, persists throughout morphogenesis, leading to neurons with non-pyramidal somata.

**Figure 5.**
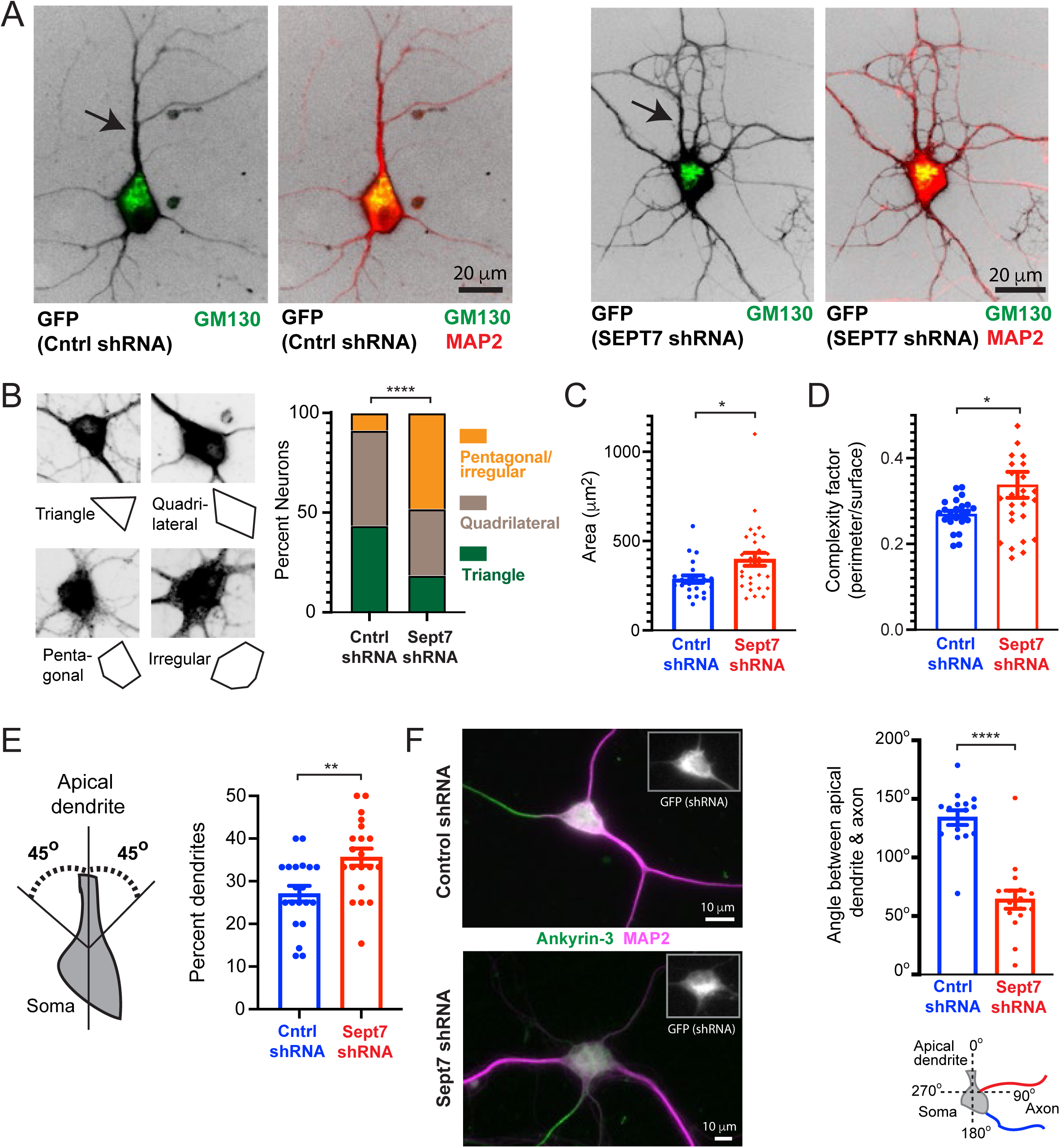
Septin-mediated control of early neuritogenesis is required for the development of a pyramidally shaped soma with proper axodendritic orientation. (A) Images of rat hippocampal neurons (DIV10), which were stained for MAP2 and GM130 after transfection with plasmids expressing GFP and scrambled control or Sept7 shRNAs at DIV1. Arrow points to the principal dendrite (apical dendrite). (B) Images show examples of GFP-expressing somata and their corresponding geometrical outlines in rat hippocampal neurons (DIV10), which were transfected with scrambled control shRNAs (triangle, quadrilateral) and Sept7 shRNAs (pentagonal, irregular) at DIV1. Stacked bar graph shows the percentage of neurons (n = 23-27) with triangular, quadrilateral or pentagonal/irregular somata. Data were analyzed with a chi-square test. (C-D) Bar graphs show the mean (± SEM) surface area and complexity factor (ratio of perimeter length to surface area) of the soma in DIV10 neurons (n = 23-28), which were transfected with shRNAs at DIV1. Data were analyzed with the Mann-Whitney U test. (E) Schematic shows the area, which is demarcated by a 45^°^ angle left and right of the principal dendrite (apical dendrite), and was used for scoring dendrites proximal to the apical dendrite. Bar graph shows the mean (± SEM) percentage of total dendrites per neuron (n = 20) within a 45° angle from the principal dendrite of neurons, which expressed shRNAs/GFP for 9 days and stained for MAP2 at DIV10. Data were analyzed with the Mann-Whitney U test. (F) Images of rat hippocampal neurons (DIV9), which were stained for MAP2 (dendrites) and ankyrin 3 (axon initial segment) after transfection with plasmids expressing GFP (inset) and scrambled control or Sept7 shRNAs at DIV1. (G) Bar graph shows the mean (± SEM) degree of the angle between the axon and principal dendrite in neurons (DIV9; n = 15-16) which were transfected with scrambled control or Sept7 shRNAs at DIV1 for 9 days. Schematic shows axon positioning with respect to the principal apical dendrite in control (blue) and Sept7-depleted neurons (red). Data were analyzed with the Mann-Whitney U test. Statistics. *p<0.05, **p<0.01, ****p<0.0001, ns: non-significant

Given that the pyramidal soma generates a dendritic tree with a principal dendrite (apical dendrite), which extends from the pyramidal apex, and basal dendrites that originate from the pyramidal base, we analyzed the morphology and orientation of dendrites. In two-dimensional neurons, the presumptive apical dendrite is the principal dendrite, which is characterized by a wider shaft and presence of the Golgi complex at the base of its shaft (Figure 5A, arrow) (Wu, et al., 2015). By identifying the presumptive apical dendrite, we analyzed the spatial distribution of dendrites and position of the axon with respect to the apical dendrite. Because in Sept7-depleted neurons, more dendrites appeared to localize near the principal apical dendrite (Figure 5A), we measured the number of dendrites as the percentage of total dendrites localizing within a 45° angle clockwise and counter-clockwise of the principal dendrite (Figure 5E). Sept7 knock-down resulted in a ∼30% increase in the percentage of dendrites, which are positioned within a 45° angle of the principal dendrite (Figure 5E). Thus, Sept7-depletion alters the spatial distribution of the dendritic tree.

We next examined whether Sept7 depletion impacts axon development and positioning. We quantified axon specification by analyzing the formation of a tau-1 positive neurite which has twice the length of any other neurite at 48, 72 and 96 h after transfection of DIV1 neurons with shRNAs. Although Sept7 depletion delayed axon specification, all neurons developed an axon after 96 h (DIV5). However, and as previously reported, axons were shorter (Ageta-Ishihara, et al., 2013; Hu, et al., 2012). In addition, axons were positioned closer to the apical dendrite (Figure 5F). Quantification of the angle of axon orientation in neurons (DIV10) showed that the axon was oriented closer to the apical dendrite of Sept7-depleted neurons. Axons originated from the soma at a 64° angle from the apical dendrite, which was ∼50% narrower than the axon-dendrite angle (134°) in control neurons. Collectively, these data show that in Sept7-depleted hippocampal neurons, defective neuritogenesis impairs the development of pyramidally shaped somata and the proper orientation of their axodendritic processes.

The phenotypes of Sept7 depletion in cultured hippocampal neurons indicates that Sept7 plays an important role in the early stages of neuronal morphogenesis in the developing brain. Because *Sept7* knock-out causes early embryonic lethality in mice (Menon, et al., 2014), we sought to test whether Sept7 depletion affects the development of cortical pyramidal neurons by knocking down Sept7 expression in mouse embryos. Using in utero electroporation, we delivered shRNAs into the layer II/III of the developing neocortex of murine E15.5 embryos, and analyzed the effects after seven days in the cortex of P3 animals. At this stage, cortical pyramidal neurons have migrated from the ventricular zone to the cortical plate (CP), and developed apical dendrites and axons, which are oriented toward and away from the CP, respectively (Figure 6A-B). Consistent with the phenotype of enlarged somata in cultured hippocampal neurons, we found Sept7 depletion altered the in vivo morphology of mouse cortical neurons, which exhibited enlarged lamellipodia- and dactylopodia-like protrusions (Figure 6B; see arrowheads in regions I and II). The mean surface area of these neurons was markedly increased (Figure 6C), and their shape was more contorted and irregular in comparison to control neurons which had primarily triangular or quadrilateral shapes (Figure 6D). In Sept7-depleted neurons, apical dendrites often emanated from wider lamellar extensions, which increased their width (Figure 6E), and were characterized by a higher incidence of kinks and bends (Figure 6F; see also arrow in Figure 6, region III). In addition to these morphological phenotypes, a number of Sept7-depleted neurons were abnormally oriented, assuming positions that were more in parallel than vertical to the CP (Figure 6B, see arrow in region I). We quantified this phenotype by measuring the angle between the axis, which is orthogonal to the pial surface, and the long axis of the neuronal soma or the linear vector that is best aligned with the proximal segment of the apical process (Figure 6G). In Sept7-depleted neurons, neuronal somata and apical dendrites were less orthogonally oriented to the pial surface (Figure 6H-I). These in vivo defects in the morphology and orientation of mouse cortical neurons, which resembled the phenotypic deficits of Sept7 depletion in rat hippocampal neurons, demonstrate that the septin cytoskeleton has a conserved function in the morphogenesis of pyramidal neurons independently of species and brain region.

**Figure 6.**
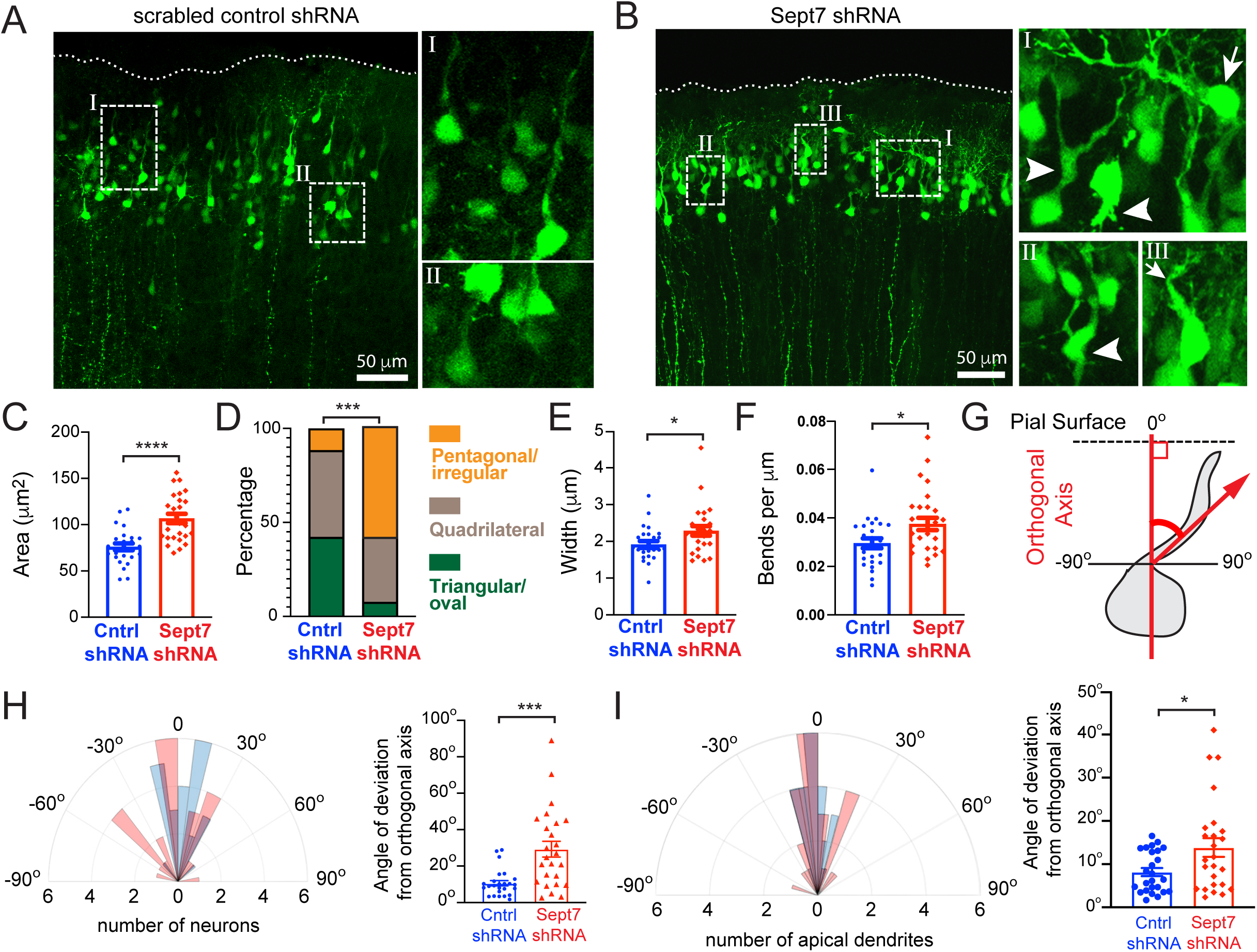
Sept7 depletion alters the morphology and orientation of pyramidal cortical neurons in the developing mouse neocortex. (A-B) Images show GFP, a marker of shRNA expression, in cortical brain sections of mice (P3) after in utero electroporation of plasmids encoding for GFP and scrambled control (A) or Sept7 shRNAs (B) into layer II/III of E15.5 embryo brains. Selected regions (I, II, III) are shown in higher magnification. Arrowheads (B, regions I and II) point to enlarged somata with extended lamellipodia-like protrusions. Pial surfaces are outlined with dotted lines. Arrows point to a neuron with its soma oriented parallel to the pial surface (B, region I) and with a bent in its apical dendrites (B, region III). Scale bar, 50 μm. (C-D) Bar graphs show the mean (± SEM) surface area of the soma (C) and the percentage of distribution of neurons (D) with somata of triangular/oval, quadrilateral, and pentagonal or irregular shapes. Analysis was performed in neurons (n = 26) from the brains of three P3 mice, which were in utero electroporated (E15.5) with scrambled control or Sept7 shRNAs. Statistical analysis was performed with an unpaired t-test (C) and chi-squared test (D). (E-F) Bar graphs show the mean (± SEM) width of apical dendrite (E; n = 26) and number of apical dendrite bents per μm length in neurons (F; n = 25) from brains of P3 mice, which were transfected with scrambled control or Sept7 shRNAs at E15.5. Quantifications were performed from three different mice and data were analyzed with the Mann-Whitney U-test (E) and unpaired t-test (F). (G) Schematic depicts the quantification of the orientation of the apical dendritic process with respect to the orthogonal apicobasal axis of the mouse neocortex. Using the horizontal axis of the pial surface as a reference, an orthogonal line (red) was drawn to define the 0° degree point from which the angle with the axis of the apical dendrite is measured. Orientation of the soma was similarly quantified by measuring the angle between the long axis of the soma and the orthogonal axis of the neocortex. (H-I) Polar histograms show the distribution of the orientation of the long axis of neuronal somata (H) and apical dendrites (I) with respect to the orthogonal axis of the neocortex – angle between the long axis of the soma and the apicobasal orthogonal axis. Negative and positive values correspond to angles left and right of the orthogonal axis, respectively. Bar graphs show the mean (± SEM) angle of deviation from the orthogonal axis irrespective of left/right orientation. Quantifications were performed from the brains of three different mice (n = 25 neurons). Data were analyzed with an unpaired t-test. Statistics. *p<0.05, ***p<0.001, ****p<0.001

## Discussion

Neuronal morphogenesis and the establishment of neuronal circuits begins with the formation of neurite processes and their differentiation into axons and dendrites. Elucidating the mechanisms of neurite initiation is key for understanding and treating a variety of neurodevelopmental and mental health disorders (e.g, autism, schizophrenia), which arise from aberrations in neurite abundance, morphology and distribution (Liu and Jan, 2020; Prem, et al., 2020; Copf, 2016; Kulkarni and Firestein, 2012). Despite advances in the formation of filopodia, which are the precursors of neurites, it is poorly understood how neuronal somata develop a pyramidally shaped organization with neurites that emerge from the vertices of narrowing cell edges.

Spatial control and tuning of the contractile and protrusive activities of the actin cytoskeleton is of key importance to neuritogenesis. In early stages of neurite formation, the yin and yang of filopodia and lamellipodia is evident by the shift of balance between these two protrusion types in phenotypes of proteins with roles in actin polymerization and organization (Sainath and Gallo, 2015; Flynn, 2013). It is poorly understood, however, how filopodial and lamellipodial activities are balanced - a problem that is compounded by filopodia forming within the actin networks of lamellipodia (Yang and Svitkina, 2011). Suppression of protrusive activity along the length of growing neurites, and a competition between myosin II and Arp2/3 activity in growth cones suggests that protrusion and contractility are similarly balanced in the soma during neurite budding (Yang, et al., 2012; Mingorance-Le Meur and O’Connor, 2009; Lin, et al., 1996; Lin and Forscher, 1995). Here, we discovered a novel cytoskeletal network, a circumferential septin wreath-like meshwork, which controls contractility and protrusion in the soma by differentially regulating the localization of myosin II and Arp2/3 at the base of nascent neurites.

Septins are essential components of neuronal morphogenesis with roles in axodendritic development and the formation of axon branches, and dendritic spines (Radler and Spiliotis, 2022; Tada, et al., 2007). In differentiating neural crest cells, septins determine the cortical sites of neurite re-emergence after mitosis, but how septins function in neurite initiation is not understood (Boubakar, et al., 2017). In pyramidal neurons, previous work showed that Sept7 depletion stunts axodendritic growth and arborization, but Sept7 was targeted after neurite formation (DIV4, stage 3 neurons) (Ageta-Ishihara, et al., 2013; Tada, et al., 2007; Xie, et al., 2007). In our studies of stage 1 (DIV0) neurons, we found a filamentous Sept5/7/11 network, which is the first of its kind in overall organization and position with respect to the microtubule and actin networks. Positionally, it bears some similarity to septins localizing along the contractile transverse actin arcs of contact-naïve and migrating epithelia (Dolat, et al., 2014). Preferential association of septins with domains micron-scale membrane curvature (e.g., base-neck of axonal filopodia and dendritic spines) raises the possibility that the wreath-like network might be in part supported by the nuclear membrane or alternatively the endoplasmic reticulum (Bridges, et al., 2016; Hu, et al., 2012; Tada, et al., 2007; Xie, et al., 2007).

Septins have evolutionarily conserved functions as scaffolds and barriers that selectively regulate the localization of a diversity of proteins (Spiliotis and Nakos, 2021; Spiliotis and McMurray, 2020). In the early stages of neurite formation, the septin network demarcates and overlaps with a meshwork of myosin IIB filaments and puncta, which are tightly interwoven with septins. Our results indicate that the circumferential septin network provides a scaffolding function for the localization of myosin IIB at the base of filopodia, which is in agreement with previous findings of Sept7 and Sept2 interaction with the myosin II heavy chain (Wasik, et al., 2017; Joo, et al., 2007). In Sept7-depeleted neurons, reduction of myosin II from the base of filopodia is also consistent with our observation of diminished retrograde actin flow and filopodia stability (Alieva, et al., 2019; Medeiros, et al., 2006; Lin, et al., 1996). The latter might be due to effects on the crosslinking/bundling of filopodial actin filaments and/or the adhesion of filopodia with extracellular matrix (ECM), which could result from a defective actin clutch at focal adhesions due to reduced myosin II-dependent actin flow and mechanotransduction (Alieva, et al., 2019; Case and Waterman, 2015).

In Sept7-depleted neurons, reduction of myosin II from the base of filopodial actin could impact the localization and function of other actin-binding proteins such as cofilin, which associates with the proximal ends of filopodia and competes with myosin II for actin binding (Hylton, et al., 2022; Elam, et al., 2013; Wiggan, et al., 2012). Loss of myosin II may increase the levels of cofilin at the base of filopodia, which could be responsible for the higher incidence of neurite bending or buckling at the cofilactin/actin boundary (Hylton, et al., 2022; Suarez, et al., 2011). In migrating cells, Sept7 over-expression correlates with increased levels of phosphorylated cofilin, suggesting that Sept7 knock-down may reduce the phosphorylation of cofilin and enhance its severing activity (Hou, et al., 2016). However, F-actin severing by cofilin promotes neuritogenesis by enabling retrograde actin flow and microtubule protrusion (Flynn, et al., 2012).

The circumferential septin network appears to suppress Arp2/3 localization and activity at the base of filopodia. To our knowledge, there is no precedence of direct inhibition of Arp2/3 activity or actin-binding by septins. In the axonal filopodia of sensory neurons, Sept7 functions downstream of the Arp2/3-mediated transition of actin patches to filopodia, which is aided by recruitment of cortactin by Sept6 (Hu, et al., 2012). We were unable to detect Sept6 at the Sept5/7/11 circumferential network, which is likely due to Sept11 being the predominate subunit of the Sept6 family and therefore, taking the position of Sept6 in the septin protomers (Cavini, et al., 2021). We posit that Sept7 functions differently in complex with Sept5/11 in murine pyramidal neurons than its previous role with Sept6 in chick dorsal root ganglia. It is plausible that Sept5/7/11 complexes compete with Arp2/3 for binding to actin filaments and/or inhibit the binding of Arp2/3 activators. Alternatively, Sept7 may indirectly suppress Arp2/3 through myosin II as recently proposed for the formation of dactylopodia in endothelial cells (Figueiredo, et al., 2021). In this scenario, downregulation of Arp2/3 would be the result of feedback loop inhibition of the β-PIX/Rac pathway, which is triggered by an upregulation of focal adhesion maturation due to the enhanced mechanotransduction by myosin II at the base of filopodia (Figueiredo, et al., 2021; Alieva, et al., 2019).

Neurite morphogenesis is accompanied by microtubule sliding and entry into membrane protrusions (Miller and Suter, 2018; Flynn, 2013; Lu, et al., 2013). In sensory neurons, Sept7 promotes microtubule entry into axonal filopodia (Hu, et al., 2012), and in pyramidal neurons, Sept7 scaffolds the α-tubulin deacetylase HDAC6 controlling the levels of microtubule acetylation, which impacts axodendritic growth (Ageta-Ishihara, et al., 2013). The circumferential septin network localizes at the periphery of the microtubule network of the soma, and overlaps only with “pioneering microtubules” that enter into the peripheral lamellae. Although we cannot exclude the possibility that Sept5/7/11 has a role in microtubule targeting to nascent neurites, our findings are most consistent with a function in the consolidation of the soma through suppression of Arp2/3 and lamellipodial activity. Notably, this early control of protrusive activity is a crucial step not only for the development of a pyramidally shaped soma, but also for the spatial orientation of neurites. Sept7 depletion altered the shape and size of the somata, and the orientation of their dendritic processes in rat hippocampal neurons in vitro and mouse cortical neurons in vivo. Therefore, septin-mediated control of early neuritogenesis provides an important function for the morphogenesis of pyramidal neurons, which is conserved in different neuronal types and species.

In sum, the somata of pyramidal neurons contain a hitherto unknown cytoskeletal network of septins (Sept5/7/11), which controls the balance of lamellipodial and filopodial protrusions during neurite initiation – a crucial morphogenetic step in the development of pyramidal neurons. Our results have implications for the pathology of autism spectrum and mental health disorders. Sept7 is directly phosphorylated by the thousand and one amino acid protein kinase 2 (TAOK2), an autism susceptibility and schizophrenia risk factor (Yadav, et al., 2017). Sept5 maps to the 22q11.2, a chromosomal locus with deletions in autism spectrum disorders and schizophrenia, and both SEPT5 and SEPT11 are upregulated in the brains of patients with schizophrenia and bipolar disorder (Harper, et al., 2012; Suzuki, et al., 2009; Pennington, et al., 2008). Thus, aberrations in the septin network and its functions may underlie the developmental deficits of neuritogenesis. Future work will explore this link, offering new insights into the pathogenesis and treatment of autism spectrum disorders and/or schizophrenia.

## STAR Methods

### Key resources table

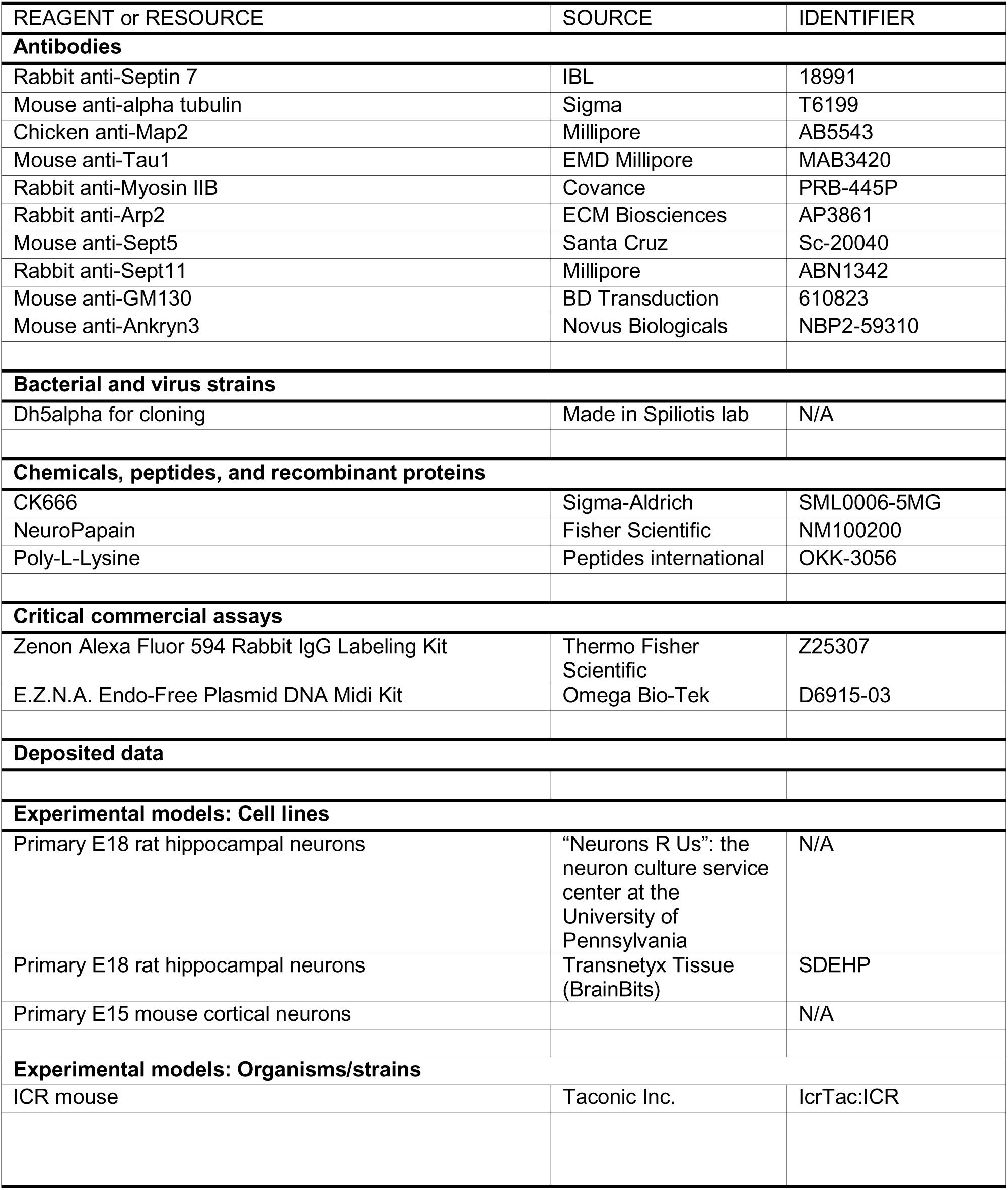

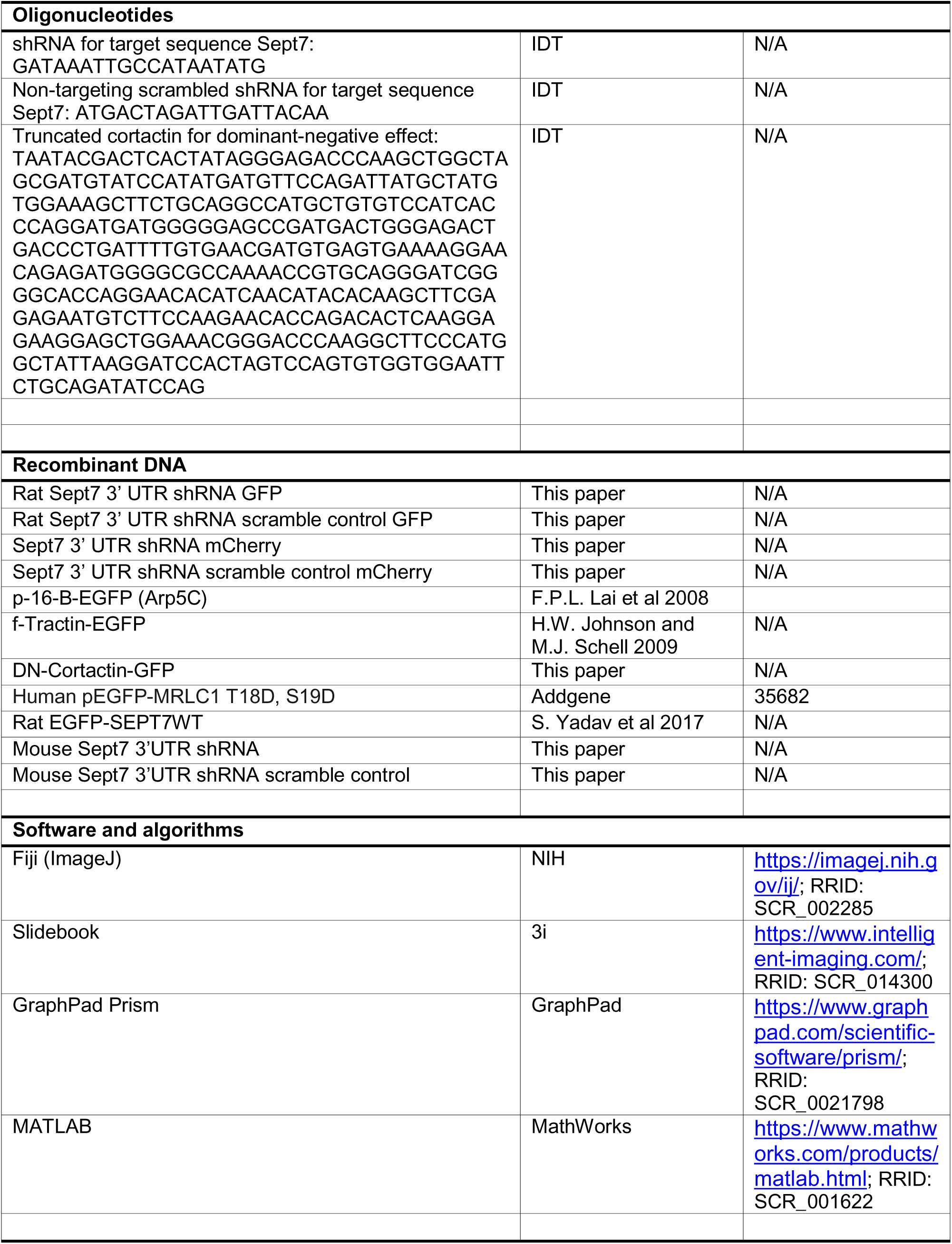

### Resource availability

#### Lead Contact

Requests for further information, resources and/or reagents should be made to the lead contact: Dr. Elias Spiliotis

#### Materials Availability

This study did not generate any new reagents.

### Experimental models and subject details

#### Primary neuronal cultures and mouse lines

Primary rat embryonic hippocampal neurons were derived from hippocampi that were isolated from the brains of mixed sex embryos, which were taken from timed pregnant Sprague-Dawley rats at 18 days of gestation. Fresh hippocampi were purchased from Transnetyx Tissue (BrainBits) or neurons were purchased in culture from the Neuron Culture Service Center (University of Pennsylvania). Neurons from hippocampi were whole hippocampi were obtained using 2 mg/mL Neuropapain in Hibernate minus Calcium media (Transnetyx Tissue) according to BrainBits neuron dissociation protocol. Neurons were cultured on surfaces coated with 1 mg/mL poly-L-Lysine (Peptides International) in Neurobasal medium supplemented with 2% B27 (ThermoFischer), and maintained at 37°C in a incubator supplemented with 5% CO_2_. Neurons were plated on 12 mm round glass coverslips (Belco Glass), which were placed in 24-well polystyrene dishes (Corning), at a density of 60,000 cells per well. For time-lapse microscopy, neurons were plated on 35 mm polystyrene dishes (VWR) at a density of 0.75-1.5 million cells per dish.

Primary mouse cortical neurons were isolated at stage E15 as previously described (Liu, et al., 2021). Following euthanasia of the dam, the embryos were quickly removed from the pregnant mouse. Embryos were then transferred to 1x Ca2+/Mg2+-free Dulbecco’s PBS (Cenesee Sci., D-PBS). Dissection of the cerebral cortex was followed by treatment with 0.01% Trypsin in D-PBS for 5 minutes at room temperature. The trypsin was then inactivated by adding 100 µm of 50 mg/ml bovine serum albumin. Dissociated neurons were seeded onto coverslips coated with 100ng/ml poly-D-lysine and 100 ng/ml laminin in NeuroBasal medium containing 1% penicillin/streptomycin (Corning), 1% GlutaMAX (Gibco), and 1x B27.

ICR mice were obtained from Taconic Inc, and timed pregnant mice were obtained by setting up the mating in the animal facility.

### Method Details

#### Transfection and in utero electroporation

Cultured neurons were transfected at DIV0 or DIV1 using Lipofectamine 3000 (ThermoFisher). All DNAs used for transfection were prepared using an Endotoxin-free plasmid isolation kit (Omega Bio-tek). Unless otherwise noted, cells were transfected for 48 hours before fixation or imaging. To reinitiate neurite formation, neurons were detached from their culture dish after 48 h of transfection using 0.25% Trypsin in sterile PBS, and re-plated in 4-chamber glass bottom 35 mm dishes (Cellvis) coated with 1 mg/mL poly-L-lysine at a density of 60,000 cells per well.

In utero electroporation was performed as described previously (Cornell, et al., 2016). Following anesthesia of pregnant dams, the uterine horns were exposed, and one to two microliters of plasmid (1-2 µg/µl) was injected into the lateral ventricle of the E15.5 embryo brain using a pulled-glass micropipette. Using a Nepa GENE CUY21 electroporator, three pulses of 32 V were applied to the embryonic brain using tweezers electrodes. The uterine horns were placed within the abdomen, and pups were allowed to recover and mature. P3 brains were dissected for morphological analysis.

#### Plasmids and Cloning

To deplete embryonic rat hippocampal neurons of Sept7, an shRNA was generated to target the 3’ UTR of the rat Sept7 mRNA. Primers encoding the shRNA sequence were ordered from IDT (Forward:5’- GATCCCCGGAAAGTCGACATTAATCATTCAAGAGATGATTAATGTCGACTTTCCTTTTTA-3’, Reverse: 5’- AGCTTAAAAAGGAAAGTCGACATTAATCATCTCTTGAATGATTAATGTCGACTTTCCGGG-3’) and were ligated and inserted to the p-Super EGFP backbone purchased from Addgene using the BglII and HindIII cloning sites. The rat Sept7 3’UTR shRNA scramble GFP construct was made using a scrambled version of the shRNA sequence, Forward: 5’- GATCCCCATGACTAGATTGATTACAATTCAAGAGATTGTAATCAATCTAGTCATTTTTTA-3’, Reverse: 5’- AGCTTAAAAAATGACTAGATTGATTACAATCTCTTGAATTGTAATCAATCTAGTCATGGG-3’. The mCherry version of these rat Sept7 3’UTR shRNA constructs was made by removing the GFP tag using the AgeI and BsrG1 cloning sites, and isolating and inserting the mCherry tag digested similarly into the pSuper backbone. Plasmid sequences were confirmed by sequencing (Genewiz).

For *in vivo* mouse experiments, shRNAs were cloned into the PSCV2-venus plasmid (Liu, et al., 2021; Hand and Polleux, 2011). The plasmid was digested with HindIII and BamH1, and the oligonucleotides containing the Sept7 3’ UTR shRNA targeting sequence, (Forward: 5’- GATCCGATAAATTGCCATAATATGTTCAAGAGACATATTATGGCAATTTATCTTTTTGGAAA-3’, Reverse: 5’- AGCTTTTCCAAAAAGATAAATTGCCATAATATGTCTCTTGAACATATTATGGCAATTTATCG-3’), or the scramble targeting sequence, (Forward: 5’- GATCCATGACTAGATTGATTACAATTCAAGAGATTGTAATCAATCTAGTCATTTTTTGGAAA-3’, Reverse: 5’- AGCTTTTCCAAAAAATGACTAGATTGATTACAATCTCTTGAATTGTAATCAATCTAGTCATG-3’) were synthesized and purchased by Integrated DNA Technologies. ShRNA oligos were annealed and then ligated to the digested pSCV2-venus plasmid. Plasmids purified with a mini-prep kit (Zippy Plasmid Miniprep, Zymo Research) and were evaluated by sequencing (ASENTA).

The plasmid encoding ArpC5B-EGFP was a kind gift from Dr. Klemens Rottner, and was previously made as previously described (Lai, et al., 2008).

The plasmid encoding EGFP-F-Tractin (EGFP-ITPKA) was a kind gift from Dr. Tanya Svitkina and was created as described previously (Johnson and Schell, 2009).

The plasmid encoding a dominant negative cortactin, DN-Cortactin-HA (Weed, et al., 2000), was created using only the N-terminal acidic domain sequence as a G-Block (IDT) with the sequence: 5’- TAATACGACTCACTATAGGGAGACCCAAGCTGGCTAGCGATGTATCCATATGATGTTCCAGATTATGCTATGTGGAAAGCTTCTGCAGGCCATGCTGTGTCCATCACCCAGGATGATGGGGGAGCCGATGACTGGGAGACTGACCCTGATTTTGTGAACGATGTGAGTGAAAAGGAACAGAGATGGGGCGCCAAAACCGTGCAGGGATCGGGGCACCAGGAACACATCAACATACACAAG CTTCGAGAGAATGTCTTCCAAGAACACCAGACACTCAAGGAGAAGGAGCTGGAAACGGGACCCAAGGCTTCCCATGGCTATTAAGGATCCACTAGTCCAGTGTGGTGGAATTCTGCAGATA TCCAG -3’ into pcDNA3.1(+) IRES GFP (Addgene) by digesting the backbone using NheI and BamHI.

The plasmid encoding human constitutively active myosin regulatory light chain (pEGFP-MRLC1 T18D, S19D) was purchased from Addgene.

Rat EGFP-Sept7WT was a kind gift of Smita Yadav, and was made as previously described (Yadav, et al., 2017).

#### Fixation and staining

Rat hippocampal neurons were fixed for 10 minutes with PBS buffer containing 4% paraformaldehyde (PFA) and 4% sucrose, permeabilized with GDB (30 mM sodium phosphate pH 7.4, 0.2% gelatin, 450 mM NaCI) containing 0.01-0.05% Triton X-100 for 10 min, and blocked with GDB for an additional 20 minutes. Primary antibodies were diluted in GDB and spun at 50,000xg for 10 min at 4°C before adding on neurons. Primary antibodies were incubated at 4°C overnight. Secondary antibodies were also diluted in GDB, spun at 50,000xg before adding to neurons for 1 h at room temperature. Samples were mounted with FluorSave hard mounting medium (EMD Millipore). Mouse cortical neurons were fixed after 2 hours in culture with 4% paraformaldehyde at room temperature for 15 minutes, and washed with PBS. Subsequently, they were permeabilized and stained as aforementioned for the rat hippocampal neurons.

Electroporated mouse brains were fixed in 4% PFA in PBS overnight at 4°C, then cryoprotected in 25% sucrose in PBS for 48 hours at 4°C. The brains were embedded with an O.C.T. compound (Sakura). Cryosections of 60 μm thickness were cut using a cryostat (Microm HM 505N) and air-dried. The sections were washed three times in Tris-buffered saline, and stained with 600 nM 4’, 6-diamidino-2-phenylindole, dihydrochloride (DAPI). The sections were mounted on glass slides using 90% glycerol in PBS.

#### Antibodies and Reagents

See also the Keynote Resources Table. Neurons were immunostained with the following antibodies: mouse anti-α-tubulin (DM1α, 1:500; SIGMA), rabbit anti-SEPT7 (1:500; IBL America), rabbit anti-Myosin IIB (1:500, Covance), rabbit anti-Arp2 (1:100, ECM Biosciences), chicken anti-MAP2 (1:2000; EMD Millipore), mouse anti-Tau1 (1:500, EMD Millipore), mouse anti-GM130 (Golgi, 1:200, BD Transduction), mouse anti-Ankryn 3 (1:100, Novus Biologicals), mouse anti-SEPT5 (1:500, Santa Cruz), rabbit anti-SEPT11 (1:500, Millipore). F(ab’)_2_ fragment affinity-purified secondary antibodies (1:200) were purchased from Jackson ImmunoResearch Laboratories and included donkey anti-mouse, -rabbit, and -chicken antibodies conjugated with AMCA, Alexa488, Alexa594 or Alexa647. To co-stain for Sept7 and MyoIIB and Arp2, rabbit anti-MyoIIB or Arp2 primary antibody was conjugated with anti-rabbit Alexa Fluor 594 using the Zenon Rabbit IgG Labeling Kit (Thermo Fisher Scientific). To stain actin, phalloidin conjugated with iFluor 647 (1:200, Abcam) was used.

#### Microscopy

Super resolution 3D structured illumination microscopy (SIM) imaging was performed with the OMX V4 microscope (GE Healthcare) using a 60X/1.42 NA objective, a z-step size of 0.125 μm and immersion oil with refractive index 1.514 (GE Healthcare).

Total internal reflection fluorescence (TIRF) imaging (20-60 frames per min) of live neurons was performed at 37°C using the TIRF module on the DeltaVision OMX V4 inverted microscope equipped with an Olympus 60x/1.49 objective and a temperature-controlled stage-top incubator. Images were acquired with sCMOS pco.edge cameras (PCO) and reconstructed with softWoRx software (Applied Precision).

Unless otherwise specified, neuronal images were obtained with a wide-field Zeiss AxioObserver Z1 inverted microscope equipped with a Zeiss 20X/0.8 dry objective, a 40X/1.2 NA water objective, a 63x/1.4 NA oil objective, a Hamamatsu Orca-R2 CCD camera and the Slidebook 6.0 software.

Overnight phase contrast or differential interference contrast microscopy (Figure 2J-N) was performed at 37°C, 5% CO2 using the Zeiss LSM700 or the Leica Stellaris 5 confocal microscopes, respectively.

Mouse brain sections (Figure 6) were imaged using the Leica TCS SP8 confocal. The images were taken with a zoom in of 1.5X and a 20X objective. The 40-80 stacks were taken at an interval of 1.04 microns in the z-direction.

#### Time-Lapse Imaging

Hippocampal neurons were transfected at DIV0 for 48 hours, trypsinized, and re-plated at a density of 60,000 cells per well on 4-chamber glass bottom 35 mm cell culture dishes (Cellvis) containing phenol red-free neurobasal media supplemented with 2% B27 (Invitrogen). Samples were placed on the microscope within 1 hour of plating and imaged for up to 12 hours to capture early neurite formation events.

For time-lapse TIRF microscopy (Figures 2H and 3E-I), neuronal medium was supplemented with 30 mM HEPES, and dishes were sealed using parafilm. Cells were imaged at 20 frames per second for 5 minutes.

For overnight phase contrast and DIC imaging, samples were kept in an imaging chamber set to 37°C, 5% CO2. Neurons transfected with Sept7 shRNA or scrambled control were identified by fluorescence microscopy, and transfected cells were imaged using phase contrast (LSM700) or DIC (Leica Stellaris) at 5-minute intervals for 12 hours.

### Quantification and statistical analysis

#### Neuronal soma quantifications

The area of neuronal somata was meausured in Fiji using the polygon selection tool, and manually drawing along the perimeter of the neuronal cell body. Selection of the cell body was guided by expression of GFP or mCherry fill from shRNA vectors. Cell bodies included all soma areas up to the neurite hillocks. Polygon selections were saved as ROIs, and the area and total length of the periphery was measured and recorded. Complexity factors were calculated by dividing the length of the perimeter by the total surface area. Nuclear areas were measured in Fiji using the Polygon selection tool and manually outlining nuclei guided by DAPI staining. Polygon selections were saved as ROIs, and the area was measured and recorded.

#### Neurite origin analysis and quantification

Neurite growth origin was used to determine whether neurites were growing from a tightly consolidated soma or from unconsolidated protrusions (e.g., hyperextended lamellipodia) of the soma. Neurites that originated from a consolidated soma were identified as those that emerged from smoothly curved regions of the soma, which were outlined by a compact actin (phalloidin) stain and sharp edges under phase contrast or DIC microscopy. Neurites that originated from lamellipodial protrusion or hyper-extended lamellae were identified as those that came from regions of the cell body with irregular and jagged curvatures, and protruded from cortical areas that lacked compact actin. Neurites bases were marked using the oval (compact somata) or rectangular (lamellipodial protrusions) selection tools in Fiji and saved as ROIs. The number of each type of neurite was recorded, and percentages for each were calculated based on the total neurites per cell.

#### Neurite number, width and branch quantification

Neurites were identified as any filopodia-like protrusions that originated from the neuronal soma or its lamellipodia and in fixed neurons contained microtubules. Neurites were counted by tracing them using the segmented line selection tool in Fiji and adding each neurite as an ROI. The total number of neurites per cell were counted and recorded.

Neurite branches were defined as any branch point on a neurite, primary, secondary, and tertiary branches included. Branch points were counted manually and marked using the oval selection tool in Fiji and saved as ROIs. The total number of branch points per cell were counted and recorded.

Neurite widths were measured using GFP or mCherry fill as a guide of membrane width. Using the straight line tool in Fiji, a 5 µm line was drawn from the neuronal cell body (base of the neurite hillock) distally along the neurite. Neurite width was measured using the straight line tool and drawing it orthogonally to the 5 µm marker. Width measurements were saved as ROIs, and their measurements were recorded.

#### Line scan and kymograph analyses

Line scans (Figure 1C, G) were used to generate plot profiles of the fluorescence intensity of Sept7 in relation to myosin IIB, Arp2, and actin (phalloidin) by drawing a 10 pixel-wide line from the center of the cell and saved as an ROI in the Fiji software. To export fluorescence intensity measurements, a plot profile was generated in Fiji using the Plot Profile plugin, which showed raw fluorescence intensity (measured as gray value) at increments of 0.4 microns. These measurements for generated for each channel in an image and exported to Excel. Fluorescence intensities were normalized to the maximum intensity of their respective protein by dividing each gray value by the maximum gray value. Ratio values were graphed using Graphpad/Prism 9.0.

Kymographs were generated from lamellar or lamellipodial regions (Figure 2H), and along the filopodia (Figure 3E-F) of hippocampal neurons using 1 pixel-wide lines and the Kymograph Builder plugin in the Fiji software.

#### Quantification of filopodia lifetime and buckling, and myosin IIB

Filopodia lifetime analysis (Figure 3G-H) was done manually using Fiji. Moving frame by frame, the segmented line tool was used to mark filopodia when they first appeared in movies, and saving those segmented lines as ROIs. After recording the collection of filopodia ROIs, each filopodia ROI was tracked for the entire duration of the movie (100 frames, 5 minutes), and the frame at which the filopodia quiesced through collapsing, retraction, engulfment, merging, or fanning was recorded. These frames were used to calculate the lifetime in seconds for each filopodia. Filopodia buckling rate (Figure 3I) was calculated by using the filopodia birth ROIs generated during filopodia lifetime analysis. Each filopodia that quiesced due to buckling was marked, and the percentage of filopodia that quiesce due to buckling was plotted (5 cells per condition).

Localization myosin IIB at the base of filopodia (Figure 3B) was quantified in the Fiji software by first generating ROIs of filopodial actin bundles in images of phalloidin-stained neurons. This provided the total count of filopodia per cell. Using the oval selection tool, ROIs of myosin IIB densities that overlapped with the filopodia ROIs were generated and saved. The myosin IIB ROIs, which overlapped with filopodia, were then counted and divided by the total number of filopodia ROIs to derive the percentage of myosin IIB-positive filopodia.

#### Quantification of axon specification, length and orientation

Axon specification and length (Figure S3) was quantified by first, identifying and marking axons as neurites which contained tau-1 and were at least twice as long as the length of any other neurites. The percentage of neurons with and without an axons was plotted (Figure S3A), and axon length (Figure S3B) was measured manually using the segmented line selection tool in Fiji. Selections were saved as ROIs, and the length was measured and recorded.

To determine the angle between the apical dendrite and axon (Figure 5F), DIV10 hippocampal neurons were stained with antibodies against MAP2, which was used to identify the apical dendrite, and Ankyrin 3 for identifying the axon initial segment. The apical dendrite was identified as the MAP2-positive dendrite with the widest shaft. After marking the center of the soma with the oval selection tool in Fiji, the angle tool of Fiji was used to draw a line from the distal end of the apical dendrite and the center of the soma. From there, the second vector of the angle was drawn to the distal portion of the marked AIS. This angle was saved as an ROI, measured and recorded.

#### Quantification of soma morphology and dendrite distribution in DIV10 hippocampal neurons

The soma of DIV10 neurons was outlined using the polygon selection tool in Fiji and the fluorescence of GFP, which was co-expressed with the shRNAs; GFP levels were adjusted to fully visualize the edges of the soma. Polygonal shapes were drawn over somata by drawing lines along the edges of the GFP fluorescence fill. Shapes were sorted based on the number of sides of each polygon (3, triangular; 4, quadrilateral; 5, pentagonal, >5, irregular) and somata were binned into each shape category. The percentage of each category was calculated by dividing the number somata per each category by the total number of neurons counted. The area of each soma and complexity factor were calculated as described above.

Hippocampal neurons were stained with antibodies against MAP2 and GM130 to identify apical dendrites. The apical dendrite was identified by the position of the Golgi (GM130), which localizes at the base of apical dendrites, and as the dendrite with the highest MAP2 intensity and widest proximal shaft (Wu, et al., 2015). Using the angle tool in the Fiji software, a line was drawn from the apical dendritic shaft to the center of the soma, and an area was demarcated by drawing lines at 45° angles clockwise and counterclockwise from the apical dendrite axis line (see Figure 5E). The total number of dendrites and dendrites within the demarcated area were counted and recorded. The number of dendrites within a 45° angle from the apical dendrite was calculated and divided by the total number of neurites per neuron, and calculated as percentage.

#### Quantification of cortical neuron morphology in brain slices

All analyses of in vivo data were done on layer 2/3 cortical neurons from P3 mice using confocal microscopy images, which were projected from z-stack data. The surface areas and shapes of neuronal somata (Figure 6C-D) were quantified as described above using the fluorescence fill from the transfection with the plasmids encoding for shRNA constructs. Polygonal selections were saved as ROIs, and the area was measured and recorded. Apical dendrite widths (Figure 6E) were measured by using the straight-line tool in Fiji. A 5 µm-long line was drawn from the base of the hillock of the apical dendrite into its shaft in order to define the point, at which the width of the dendrite is measured. Neurite width was measured using the straight-line tool. Width measurements were saved as ROIs, and their measurements were recorded.

An analysis of the bends per micron of the apical dendrite (Figure 6F) was conducted by measuring the length of the dendrite and counting the turns. The number of bends per micron was calculated by dividing the number of bends per apical dendrite by the length of the corresponding apical dendrite.

Apical dendrite and soma orientations were calculated as illustrated in Figure 6G. A straight line perpendicular to the pial surface of the brain slice was drawn as a reference line. Apical dendrites skewed to the right from the perpendicular line were defined as positive angles, and those skewed to the left were defined as negative angles. For statistical calculations, absolute values were used. The orientation of the soma was also measured based on its long axis using the perpendicular line as a reference as described above.

#### Fluorescence Intensity Quantifications

Neurons were identified as transfected based on the fluorescence of GFP or mCherry, which were expressed from the plasmids that encoded shRNAs. Using the GFP or mCherry fluorescence fill, neuronal areas were outlined using the segmented mask tool in Slidebook 6.0, and Mask Statistics were used to measure the sum intensity of each fluorescent channel and the overall area of the outlined neuron. The sum intensity was divided by the area to generate fluorescence intensities per area unit.

#### Statistical Analysis

Statistical analysis of data was done using GraphPad Prism 9.0 software. Each data set was tested for normal distribution of variance using the D’Agostino and Pearson, Shapiro-Wilk and Kolmogorov-Smirnov tests. For data that were normally distributed, *p*-values were derived using an unpaired t-test, if they had the same SD, or unpaired t-test with Welch correction. Data that were binned into two categories were analyzed with a Fisher’s exact test (Figure S3A) and data that were binned into more than two categories were analyzed with the chi-squared test. In Figure 6F-I, outliers were identified using the ROUT method using the ’Identify Outliers’ feature in Prism 7 (GraphPad). Data graphs were plotted in GraphPad Prism using the scatter dot plot function showing the mean and standard error of the mean (SEM). N values and p values for each experiment are denoted in the corresponding figure and/or figure legend. Statistical significance was set at 0.05.

## Acknowledgments

We thank Drs. Klemens Rottner (Helmholtz Center for Infection Research), Timothy O’Connor (University of British Columbia), Tanya Svitkina (University of Pennsylvania) and Smita Yadav (University of Washington) for constructs. We are also grateful to Drs. Harini Sreenivasappa (Drexel University) and Anthony Choo (Leica Microsystems) for help with microscopy. All microscopy was performed at Drexel University’s Cell Imaging Center. The work was supported by National Institutes of Health grants 5R35 GM136337-02 to ETS and 5R01 NS098096-05 to KT, and Commonwealth Universal Research Enhancement Formula (CURE) grants SAP 4100085747 from the Pennsylvania Department of Health to ETS and KT.

## Author Contributions

M.R.R. designed and conducted experiments and data analyses with hippocampal neurons, and contributed to the writing of the manuscript. X.L. performed experiments and data analyses with cortical neurons. M.P. and B.D. performed data analyses. K.T. designed, performed and directed in vivo experiments and analyses. E.T.S. directed the design and progress of all experiments and analyses, performed data analyses, assembled figures and wrote the manuscript.

## Declaration of Interests

The authors declare no competing interests.

## Notes

### Competing Interest Statement

The authors have declared no competing interest.

