## Supplemental Figures for "Pyramidal neuron morphogenesis requires a septin network that stabilizes filopodia and suppresses lamellipodia during neurite initiation"

SUPPLEMENTAL TEXT & FIGURES

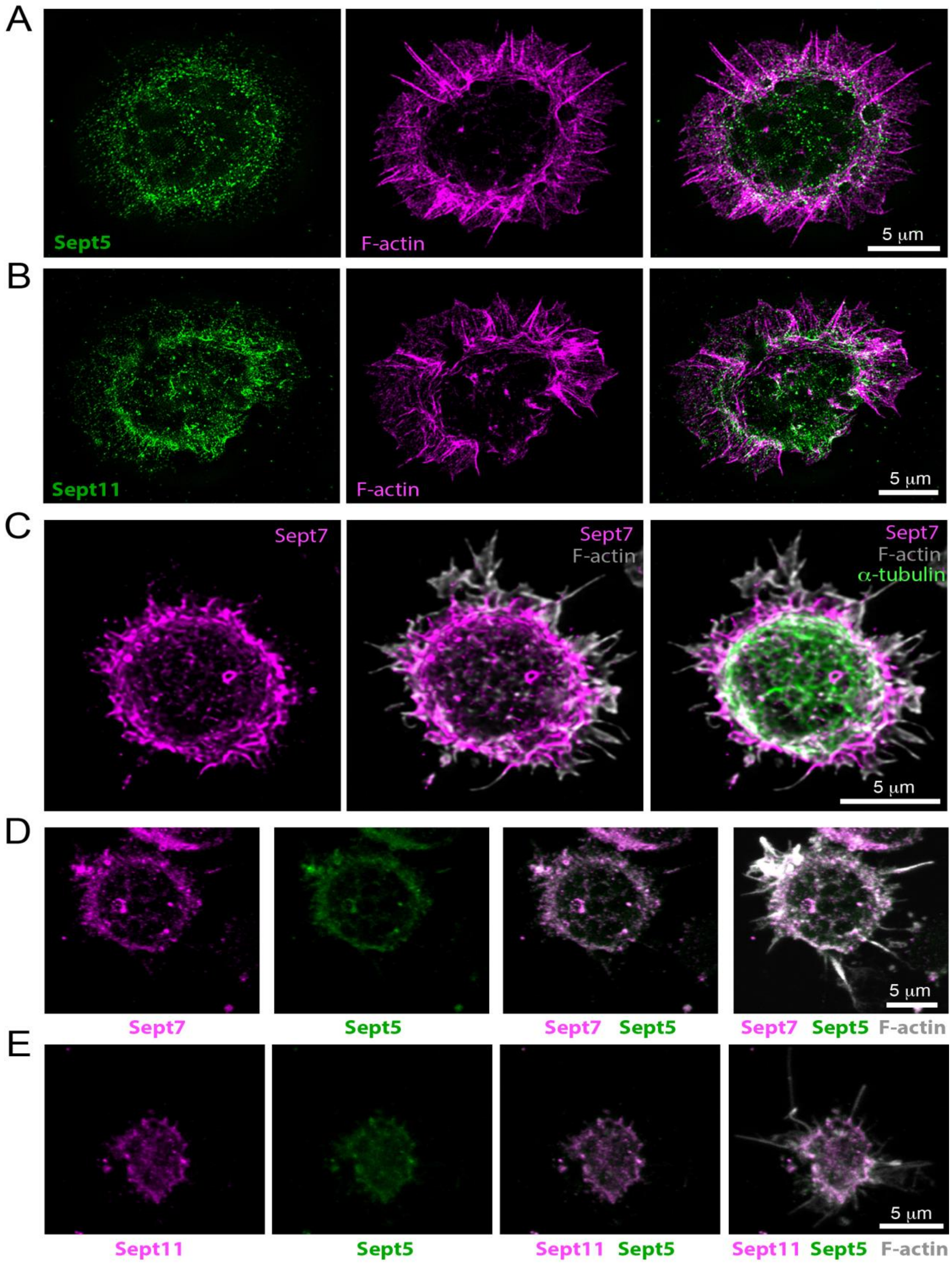


***Figure S1. Related to Figure 1.***

**Figure S1. The soma of hippocampal and cortical neurons contains a circumferential septin network that consists of septins 5, 7 and 11, related to Figure 1.**

(A-B) SIM images of embryonic (E18) rat hippocampal neurons (DIV0), which were stained with phalloidin (F-actin) and antibodies against Sept5 (A) and Sept11 (B). Scale bars, 5 mm.

(C) Confocal super-resolution (Lightning deconvolution) images of an embryonic (E15) mouse cortical neuron (DIV0), which was stained with antibodies against Sept7 and a-tubulin, and phalloidin (F-actin). Images were processed with the Lightning deconvolution

(D-E) Confocal super-resolution (Lightning deconvolution) images of embryonic mouse cortical neurons (DIV0), which were stain with phalloidin (F-actin) and antibodies against Sept7 and Sept5 (D) or Sept5 and Sept11 (E).

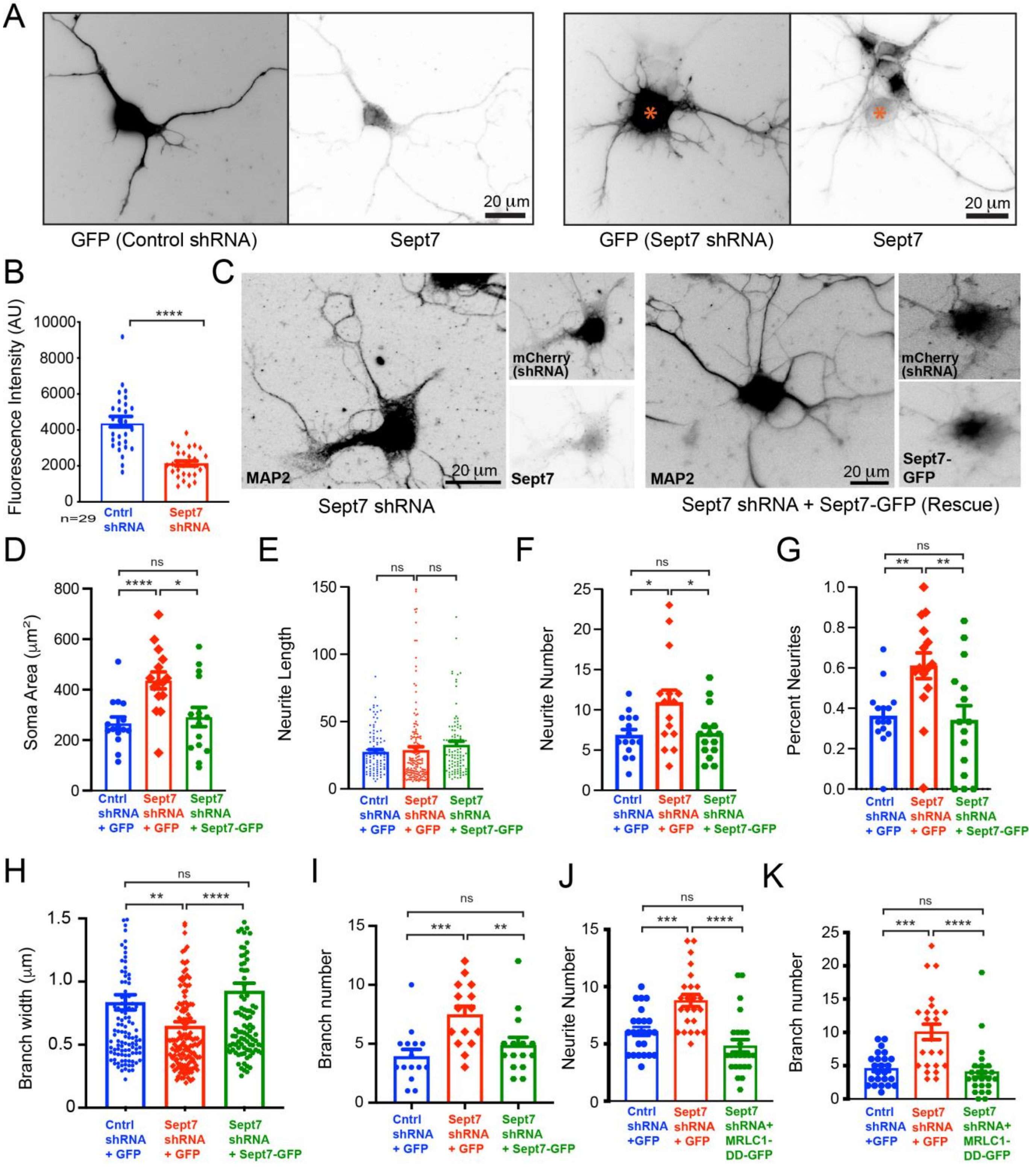


*Figure S2. Related to Figures 2 and 4.*

**Figure S2. Sept7 knock-down, and rescue of phenotypes with Sept7-GFP, related to Figures 2 and 4.**

(A) Images of rat of hippocampal neurons, which were transfected at DIV1 for 48 h with plasmids expressing GFP and scrambled control or Sept7 shRNAs, and subsequently stained with anti-Sept7. Asterisk marks the cell body of a Sept7-depleted neuron.

(B) Bar graph shows the mean (± SEM) fluorescence intensity of endogenous Sept7 in neurons (n = 29) transfected with scrambled control and Sept7 shRNAs. Data were analyzed with an unpaired t-test.

(C) Rat hippocampal neurons were transfected at DIV1 for 48 h with a plasmid expressing mCherry and shRNA against 3'UTR of *Sept7* in the absence or presence of additional plasmid expressing Sept7-GFP, which is shRNA-resistant. Neurons were stained with antibodies against MAP2 and Sept7. Images show mCherry as a reporter of shRNA expression, endogenous MAP2, Sept7-GFP or endogenous Sept7. Scale bars, 20 μm.

(D-G) Bar graphs show the mean (± SEM) surface area of the soma (D), length of neurites (E; n = 103-164, 15 neurons), neurite number per neuron (F; n = 15), and percentage of neurites that originate from lamellipodial protrusions per neuron (G; n =15). Rat hippocampal neurons (DIV1) were transfected with plasmids expressing GFP and control scrambled or Sept7 shRNAs. Rescue of Sept7 knock-down was performed by co-expression of Sept7-GFP with Sept7 shRNA and mCherry. Quantification was performed in 15 neurons and data were analyzed with the unpaired t-test and Mann-Whitney U test.

(H-I) Bar graphs show the mean (± SEM) width of neurites (H; n = 105-151, 15 neurons) and total number of neurite branches per neuron (I; n = 15). Rat hippocampal neurons (DIV1) were transfected with plasmids expressing GFP and control scrambled or Sept7 shRNAs. Rescue of Sept7 knock-down was performed by co-expression of Sept7-GFP with Sept7 shRNA and mCherry. Data were analyzed with the Mann-Whitney U test (H, I).

(J-K) Bar graphs show the mean (± SEM) number of neurites (J) and total neurite branches per neuron (K; n = 24). Quantifications were performed in rat hippocampal neurons, which were transfected that expressed control scrambled shRNA and GFP, Sept7 shRNA and GFP, and Sept7 shRNA and MRLC1-DD-GFP. Neurons that expressed shRNAs were identified by the presence of mCherry, which is encoded by the shRNA-expressing plasmid. Data were analyzed with the Mann-Whitney U test.

Statistics. *$p<0.05$, **$p<0.01$, ***$p<0.001$, ****$p<0.0001$, ns: non-significant

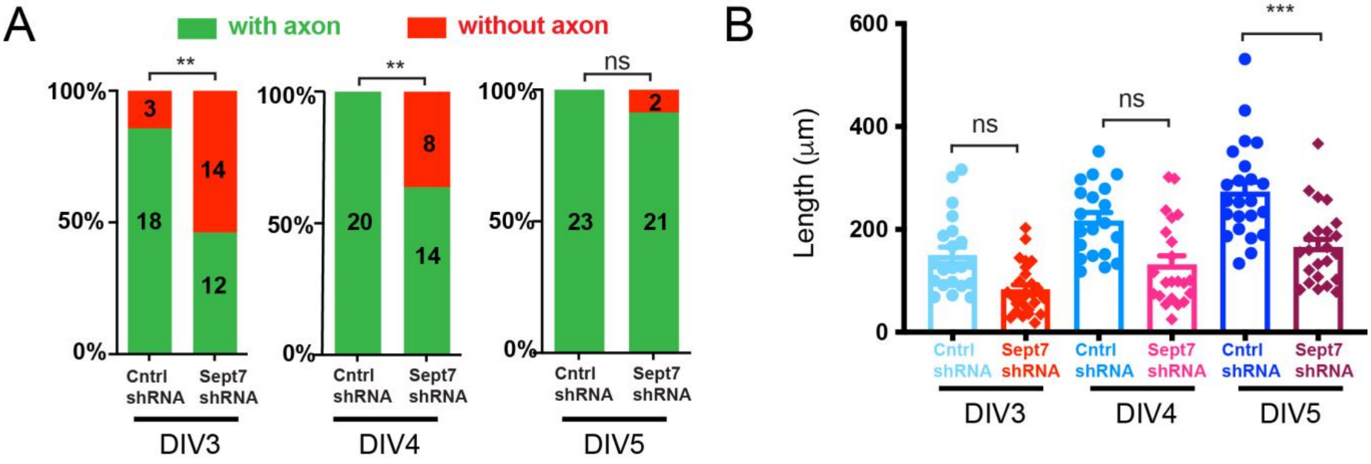


***Figure S3. Related to Figure 5.***

**Figure S3. Sept7 knock-down delays axon specification and impacts axon length, related to Figure 5.**

(A-B) Rat hippocampal neurons (DIV1) were transfected with GFP-encoding plasmid that express Sept7 or scrambled control shRNA for 48 (DIV3), 72 (DIV4) and 96 (DIV5) hours. Neurons were stained with antibodies against MAP2 and tau-1, and axonal specification was determined by the presence of a tau1-positive neurite, which was at least twice as long as any other neurite. Stacked bar graphs show percentage of neurons with (green) and without (red) axons; the number of neurons analyzed is shown in each bar. Statistical significance was derived using the Fischer's exact test.

(B) Bar graph shows the mean (± SEM) length of the axon of rat hippocampal neurons, which were transfected at DIV1 with GFP-encoding plasmid that expresses Sept7 or scrambled control shRNA for 48 (DIV3), 72 (DIV4) and 96 (DIV5) hours. Data were analyzed with an unpaired t-test.

Statistics. *$p<0.05$, **$p<0.01$, ***$p<0.001$, ****$p<0.0001$, ns: non-significant

### SUPPLEMENTAL VIDEOS

**Video S1. Retrograde actin flow and GFP-ARPC5 dynamics in control neurons, related to Figure 2.** Rat hippocampal neurons were transfected with GFP-ARPC5 and scrambled control shRNAs for 48 h, trypsinized and re-plated for 1 h before imaging with TIRF microscopy. GFP-ARPC5 is shown in inverted monochrome, and mCherry, which is co-expressed with shRNAs, is shown in cyan and fades out over time. Timer (00:00) is in minutes:seconds, and video playback rate is 20 frames per second. Scale bar, 10 µm.

**Video S2. Lamellipodia spreading and loss of retrograde actin flow in Sept7-depleted neurons, related to Figure 2.** Rat hippocampal neurons were transfected with GFP-ARPC5 and Sept7 shRNAs for 48 h, trypsinized and re-plated for 1 h before imaging with TIRF microscopy. GFP-ARPC5 is shown in inverted monochrome, and mCherry, which is co-expressed with shRNAs, is shown in cyan and fades out over time. Timer (00:00) is in minutes:seconds, and video playback rate is 20 frames per second. Scale bar, 10 µm.

**Video S3. Stable and growing neurites in control neurons, related to Figure 2.** Rat hippocampal neurons (were transfected with scrambled control shRNAs for 48 h, trypsinized, re-plated and imaged overnight with phase contrast confocal microscopy. Transfected neurons were identified by the fluorescence of GFP, which is co-expressed with shRNAs. Movie shows the neuron in Figure 2J (see inset for GFP expression). Timer is in minutes, and video playback rate is 15 frames per second. Scale bar, 10 µm.

**Video S4. Growing and shrinking neurites in control neurons, related to Figure 2.** Rat hippocampal neurons were transfected with scrambled control shRNAs for 48 h, trypsinized, re-plated and imaged overnight with phase contrast confocal microscopy. GFP, which is co-expressed with shRNAs, was used to identify transfected neurons. Movie shows the neuron in Figure 2K (see inset for GFP expression). Timer is in minutes, and video playback rate is 20 frames per second. Scale bar, 10 µm.

**Video S5. Dynamic filopodia in Sept7-depleted neurons, related to Figure 2.** Rat hippocampal neurons were transfected with Sept7 shRNAs for 48 h, trypsinized, re-plated and imaged overnight with phase contrast confocal microscopy. GFP, which is co-expressed with shRNAs, was used to identify transfected neurons. Movie shows the neuron in Figure 2L (see inset for GFP expression). Timer is in minutes, and video playback rate is 15 frames per second. Scale bar, 10 µm.

**Video S6. Filopodium-to-lamellipodium conversion, and spreading lamellipodia in Sept7-depleted neurons, related to Figure 2.** Rat hippocampal neurons were transfected with Sept7 shRNAs for 48 h, trypsinized, re-plated and imaged overnight with DIC confocal microscopy. GFP, which is co-expressed with shRNAs, was used to identify transfected neurons. Movie shows the neuron in Figure 2M (see inset for GFP expression). Timer is in minutes, and video playback rate is 20 frames per second. Scale bar, 10 µm.

**Video S7. Formation of a dactylopodium-like protrusion in Sept7-depleted neuron, related to Figure 2.** Rat hippocampal neurons were transfected with Sept7 shRNAs for 48 h, trypsinized, re-plated and imaged overnight with DIC confocal microscopy. GFP, which is co-expressed with shRNAs, was used to identify transfected neurons. Movie shows the neuron in Figure 2N (see inset for GFP expression). Timer is in minutes, and video playback rate is 20 frames per second. Scale bar, 10 µm.

**Video S8. Filopodia dynamics and engulfment by lamellipodial veils in Sept7-depleted neurons, related to Figure 3.** Rat hippocampal neurons were transfected with F-tractin-GFP and Sept7 shRNAs for 48 h, trypsinized, replated and imaged with TIRF microscopy. F-tractin-GFP is shown in inverted monochrome, and mCherry, which is co-expressed with shRNAs, is shown in cyan and fades out over time. Movie shows neuron in Figure 3E. Timer is in minutes, and video playback rate is 20 frames per second. Scale bar, 10 µm.

**Video S9. Filopodia dynamics in control neurons, related to Figure 3.** Rat hippocampal neurons were transfected with F-tractin-GFP and scrambled control shRNAs for 48 h, trypsinized, replated and imaged with TIRF microscopy. F-tractin-GFP is shown in inverted monochrome, and mCherry, which is co-expressed with shRNAs, is shown in cyan and fades out over time. Movie shows neuron in Figure 3F. Timer is in minutes, and video playback rate is 20 frames per second. Scale bar, 10 µm.
